# *In Vivo* Calcium Imaging in the Near-Infrared II Window

**DOI:** 10.1101/2025.06.26.661443

**Authors:** Danyang Xu, Zhisheng Wu, Hui Yang, Gu Chen, Wayne Jason Li, Xingyu Yue, Guang-Lei Wang, Sixin Xu, Hanze Yu, Yuanhua Liu, Zideng Dai, Haibo Jiang, Long-jiang Yu, Hongjie Dai, Feifei Wang

**Author notes:** These authors contribute equally to this work (Danyang Xu, Zhisheng Wu).

## Abstract

Non-invasive deep-tissue calcium imaging of live mammals with high sensitivity and resolution is challenging owing to light scattering experienced by traditional calcium ion (Ca^2+^) indicators with excitation and emission wavelengths within 400-750 nm. Here, we report near-infrared II (NIR-II) calcium imaging beyond 1000 nm by exploring a natural protein derived from a bacterium (*Thermochromatium tepidum*) living in a calcium carbonate-rich environment. This highly photostable fluorescent protein enables NIR-II imaging of intracellular Ca^2+^ responses to stimulant drugs in cultured mammalian cells with sensitivity comparable to that of visible Ca^2+^ indicators. We achieve *in vivo* NIR-II imaging of Ca^2+^ transients in response to two different tumor treatment strategies in intact tumors with high sensitivity, resolution, and contrast, opening the possibility of non-invasive deep-tissue calcium imaging for assessing treatment efficacy longitudinally.

**TOC Graphic:** 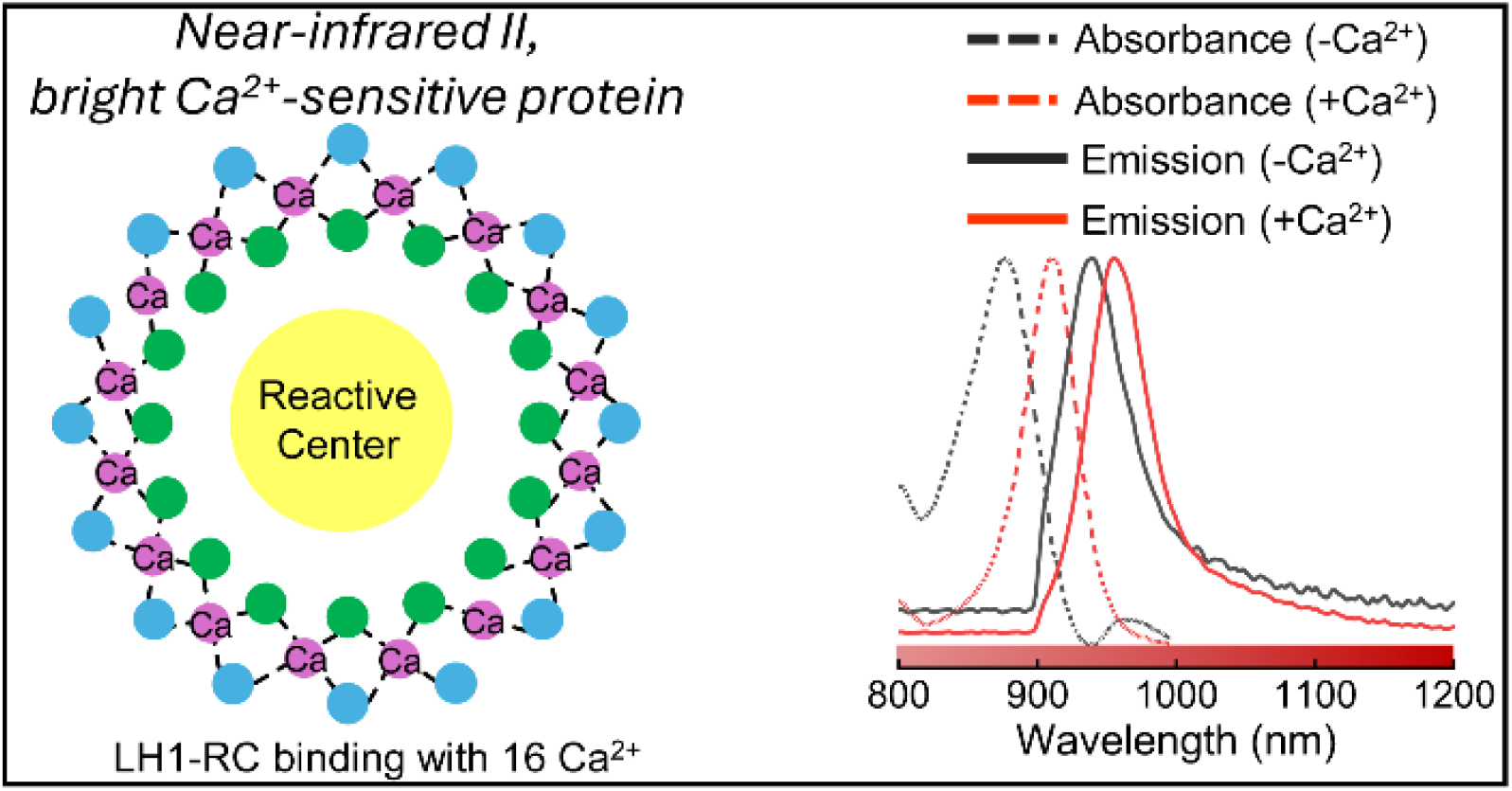

## Introduction

Calcium signaling is crucial for physiological processes^1^. Investigating calcium signals in neurons enhances the understanding of brain functions such as perception, movement, and memory^2-4^. Additionally, calcium regulates signaling pathways involved in oncogenesis and cancer progression^6^. Disruptions in calcium homeostasis connect to increased malignant phenotypes, including abnormal proliferation, resistance against cell death, migration, invasion, and metastasis^5, 6^. Dysregulated expression of calcium channels (e.g., TRP^7^, Piezo^8^) has been observed in various cancers such as breast cancer, prostate cancer, and glioma. Monitoring calcium signals in tumors offers valuable insights for clinical oncology^9^.

Visible (400-700 nm) calcium indicators, including chelators (Fluo-4^10, 11^, Fluo-8^12^) and fluorescent proteins, are widely used to detect fluctuations in intracellular^13^ or intercellular^14^ calcium ion concentrations by measuring changes in fluorescent signals. The GCaMP series is a representative visible fluorescent protein for calcium sensing^15, 16^, which has been optimized through multiple rounds of mutagenesis and selection. These enhancements include increased sensitivity and affinity to calcium ions, faster response speed, and expanded dynamic and spectral ranges^17^. Chemigenetic indicators, which combine the specificity of gene targeting with the versatility of chemical probes, offer a flexible approach to adjusting the emission wavelength for calcium sensing by appending CaM to protein scaffolds (e.g., HaloTag^18-20^). Chemigenetic calcium indicators with peak emissions at 655 nm^19^ and 765 nm^18^ have been reported. Recently, the emission peak of genetically encoded calcium ion indicators (GECI) has been extended to ∼ 700 nm, albeit at the expense of brightness, including biliverdin (BV)-based NIR-GECO1^21^ and Förster resonance energy transfer (FRET)-based iGECI ^22^. However, visible and near-infrared I (NIR-I, 700-900 nm) Ca^2+^ indicators-based imaging undergo strong light scattering, tissue autofluorescence, and limited penetration depth (∼100 µm for visible one-photon imaging)^23^ for deep tissue observation, insufficient for brain tumors and internal organ imaging and restricting their applications to cells^24, 25^, transparent animals^26^, brain slices^27^ or the superficial layer of mouse brain^28^. Two-photon or multi-photon microscopy allows for deeper tissue imaging (∼ 1 mm) with cellular resolution, but it has limitations in imaging speed and field of view^29-31^.

Fluorescence imaging in near-infrared II (NIR-II, 1000-3000 nm) or shortwave infrared (SWIR) window enables better contrast, resolution, and deeper penetration depth^32^ compared to visible and NIR-I imaging due to reduced light scattering and diminished tissue autofluorescence at longer wavelengths^33-50^. Various NIR-II molecular fluorophores and inorganic nanostructured fluorescence probes have been designed for a range of applications, including vascular and hemodynamic imaging^45^, lymph node imaging^37^, and molecular imaging^41^. The development of NIR-II Ca^2+^ indicators^51^ for deep-tissue calcium imaging is promising, but it requires high specificity and sensitivity to changes in Ca^2+^ concentrations, as well as a high quantum yield to detect fast kinetics and compensate for signal attenuation through tissues.

*Thermochromatium* (*Tch*.) *tepidum* lives in a calcium carbonate-rich environment, and its growth relies on calcium^52-57^. The photocomplex (light-harvesting 1-reaction centre complex, LH1-RC) from *Tch. tepidum* contains 16 calcium-binding sites and 32 bacteriochlorophyll *a* (BChl *a*) molecules, with its absorption exhibiting strong sensitivity and specificity to calcium^52^. Here, we present the calcium indicator derived from the natural LH1-RC of *Tch. tepidum*^52-57^, extending calcium imaging to the NIR-II window at 1000-1300 nm and demonstrating high photostability^52^. We delivered the protein into mammalian cells using a lipid encapsulation strategy and monitored the intracellular Ca^2+^ response to stimulant drugs ATP and 4-chloromethcathinone (4-CMC) in the NIR-II window. The LH1-RC enabled non-invasive *in vivo* NIR-II calcium imaging of intact tumors, facilitating real-time observation of Ca^2+^ signaling in deep tissues with high sensitivity, resolution, and contrast. The intracellular calcium signals in A549 human non-small cell lung cancer were monitored following treatment with dexamethasone and SOR-C13, providing a promising method to assess treatment efficacy.

## Results

### Fluorescent protein for Ca^2+^ sensing in the NIR-II window

Highly purified LH1-RC was isolated and collected from *Tch. tepidum*^52^ by anion exchange chromatography (Methods). The growth of this purple sulfur photosynthetic bacterium, originally found in the calcium carbonate-rich Mammoth Hot Springs in Yellowstone National Park, is significantly confined in calcium-deficient medium^58,59^. The crystal structure of LH1-RC was determined at 1.9 Å resolution^57^ (Fig. 1a; larger images observed from the side and top views can be found in Supplementary Fig. 1). LH1-RC was elliptical in shape, containing 16 pairs of light-harvesting (LH1) αβ-subunits, BChl *a*, carotenoids, and Ca^2+^ sensitive sites. The side chain of α_(n+1)_-Asp49, the carbonyl oxygens of α_(n+1)_-Trp46, α_(n+1)_-Ile51, β_n_-Trp45, and two water molecules are the ligands for Ca^2+^.The originally extracted LH1-RC exhibited a narrow absorption band at 915 nm due to calcium ions occupying the calcium-sensitive sites. After chelation with EDTA to remove Ca^2+^, the absorption peak blue-shifted to 879 nm. Subsequently, we mixed the LH1-RC with Ca^2+^ solutions at concentrations ranging from 0 to 1000 µM. The entire procedure (Supplementary Fig. 2) also demonstrated that the binding or dissociation process with Ca^2+^ is reversible. The absorption peak gradually redshifted with the increase in Ca^2+^ concentration (Fig. 1b). The presence or absence of Ca^2+^ modulates the excitonic coupling among BChl *a* molecules, which in turn alters the LH1 Qy transitions and results in corresponding changes in the absorption spectrum^60, 61^. The absorbance at 915 nm increased with the rising Ca^2+^ concentration at the µM level (Fig. 1c). The affinity constant (*K*_d_) of LH1-RC for Ca^2+^ was 63.7 μM and the Hill coefficient was 2.7. Calcium binding kinetics were assessed by monitoring changes in the fluorescence intensity of LH1-RC after the addition of Ca^2+^ to the Ca^2+^-free LH1-RC (Supplementary Fig. 3). Following the addition, the fluorescence of LH1-RC exhibited a very steep increase within the first 0.02 s, a slower rise from 0.02 to 2.38 s, and finally a relatively steady state. Comparison with other calcium indicators is summarized in Supplementary Table 2. The dynamic range of LH1-RC can be extended to 1 mM, making it suitable for calcium imaging in organelles with high calcium ion levels, such as lysosomes and the endoplasmic reticulum (ER)^62^. The increases in absorbance and fluorescence intensity induced by Ca^2+^ were significantly greater than those induced by Na^+^, Mg^2+,^ or Zn^2+^, indicating that LH1-RC has strong selectivity for calcium ions (Supplementary Fig. 4). We did not observe obvious changes in the absorbance of LH1-RC with or without Ca^2+^ in solutions with pH 6-8 (Supplementary Fig. 5).

**Figure 1.**
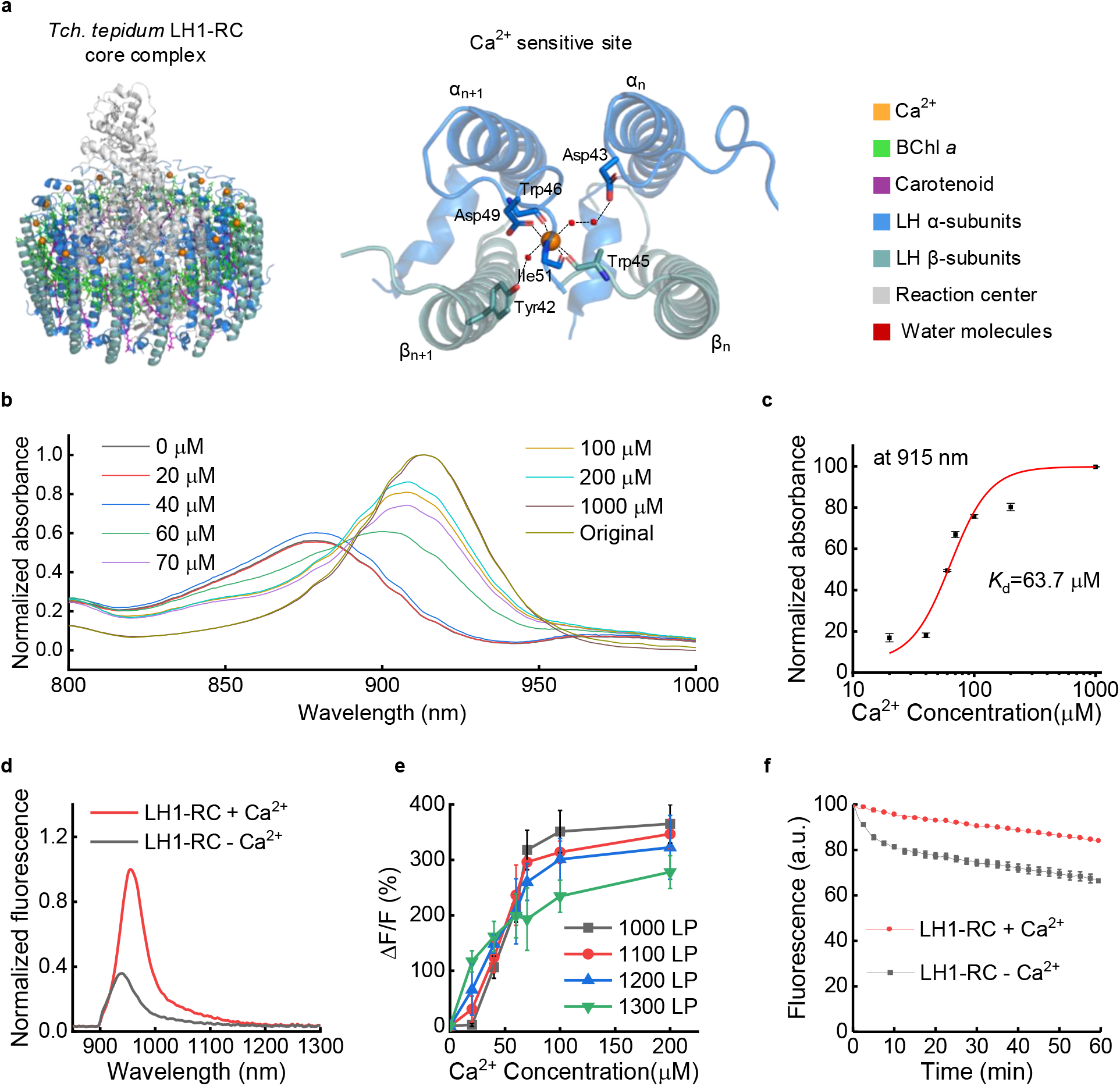
LH1-RC for NIR-II calcium imaging. (**a**) Schematic of the overall structure (viewed from the direction parallel to the membrane plane) and the Ca^2+^ sensitive sites of the LH1-RC complex. (**b**) Absorbance spectra of LH1-RC in Ca^2+^ solutions at concentrations ranging from 0 to 1000 µM. (**c**) Absorbance at 915 nm of LH1-RC in Ca^2+^ solutions at different concentrations. (**d**) Fluorescence spectra of LH1-RC with Ca^2+^ (LH1-RC + Ca^2+^) or without Ca^2+^ (LH1-RC -Ca^2+^). An 860 nm laser was used for excitation, and a 900 nm long-pass filter was applied to filter out the excitation light. (**e**) Response of LH1-RC as a function of Ca^2+^ concentration. Fluorescence was filtered using long-pass (LP) filters with cut-on wavelengths ranging from 1000 to 1300 nm. *ΔF/F= (F*_*+*Ca_*-F*_-Ca_*)/F*_-Ca_, where *F*_+Ca_ and *F*_-Ca_ represent fluorescence signals of LH1-RC with or without Ca^2+^, respectively. A 915 nm laser was used for excitation. (**f**) Photobleach curves of LH1-RC in media with or without Ca^2+^ in centrifuge tubes. A 915 nm laser with a power of 5 W was used for excitation. Fluorescence was filtered using a 1000 nm long-pass filter before being recorded by a camera. (**c,e,f**) Error bars represent s.d. for *n* = 3.

LH1-RC has an emission peak at ∼ 940 nm after EDTA treatment, which shifts to ∼ 955 nm upon binding with Ca^2+^ due to changes in the microenvironment surrounding BChls (Fig. 1d). LH1-RC with Ca^2+^ exhibits a quantum yield of 0.8% in the 1000-1400 nm range (see Methods), which is a high value compared to other NIR-II fluorophores (Supplementary Table 1), enabling imaging with an exposure time of 10 ms (Supplementary Fig. 6a). The emission tails extend into the ∼1300 nm NIR-II window. Fluorescence changes of LH1-RC in response to Ca^2+^ were characterized in different NIR-II sub-windows (Fig. 1e). A fluorescence change *ΔF/F* > 350% was observed when the Ca^2+^ concentration was greater than 70 µM, and fluorescence emission was collected within the 1000-1700 nm NIR-II window. *ΔF/F* decreased with the increase in the cut-on wavelength of long-pass filters, and fluorescence became weaker at longer wavelengths. Both LH1-RC with and without Ca^2+^ exhibited high photostability, as examined by long-term 915 nm illumination at a power of 5 W for 60 minutes (Fig. 1f). The fluorescence intensity of LH1-RC decreased by 15.9% with Ca^2+^ and 31.9% without Ca^2+^, indicating greater stability compared to previous near-infrared calcium indicators^21, 22^. LH1-RC is more stable with Ca^2+^ than without it, as binding of Ca^2+^ at the C-terminal side of LH1 polypeptides likely results in a more rigid and ordered structure than in their Ca^2+^-depleted LH1-RC^60^. The high photostability of LH1-RC facilitates long-term calcium imaging.

### NIR-II calcium imaging of mammalian cells

LH1-RC is difficult for mammalian cells to uptake directly (Supplementary Fig. 7). To enhance the delivery efficiency to C2C12 mouse myoblast cells, we encapsulated LH1-RC with Lipofectamine™ 3000 (LH1-RC@lipid, Methods). Transmission electron microscopy (TEM) of LH1-RC and LH1-RC@lipid revealed that the LH1-RC was successfully encapsulated within the lipid sphere with an average diameter of ∼ 932 nm (Fig. 2a, Supplementary Fig. 8). Dynamic light scattering (DLS) measurements showed that the average hydrated size of LH1-RC@lipid was ∼888.0 nm, consistent with the TEM results (Fig. 2b, Supplementary Fig. 8).

**Figure 2.**
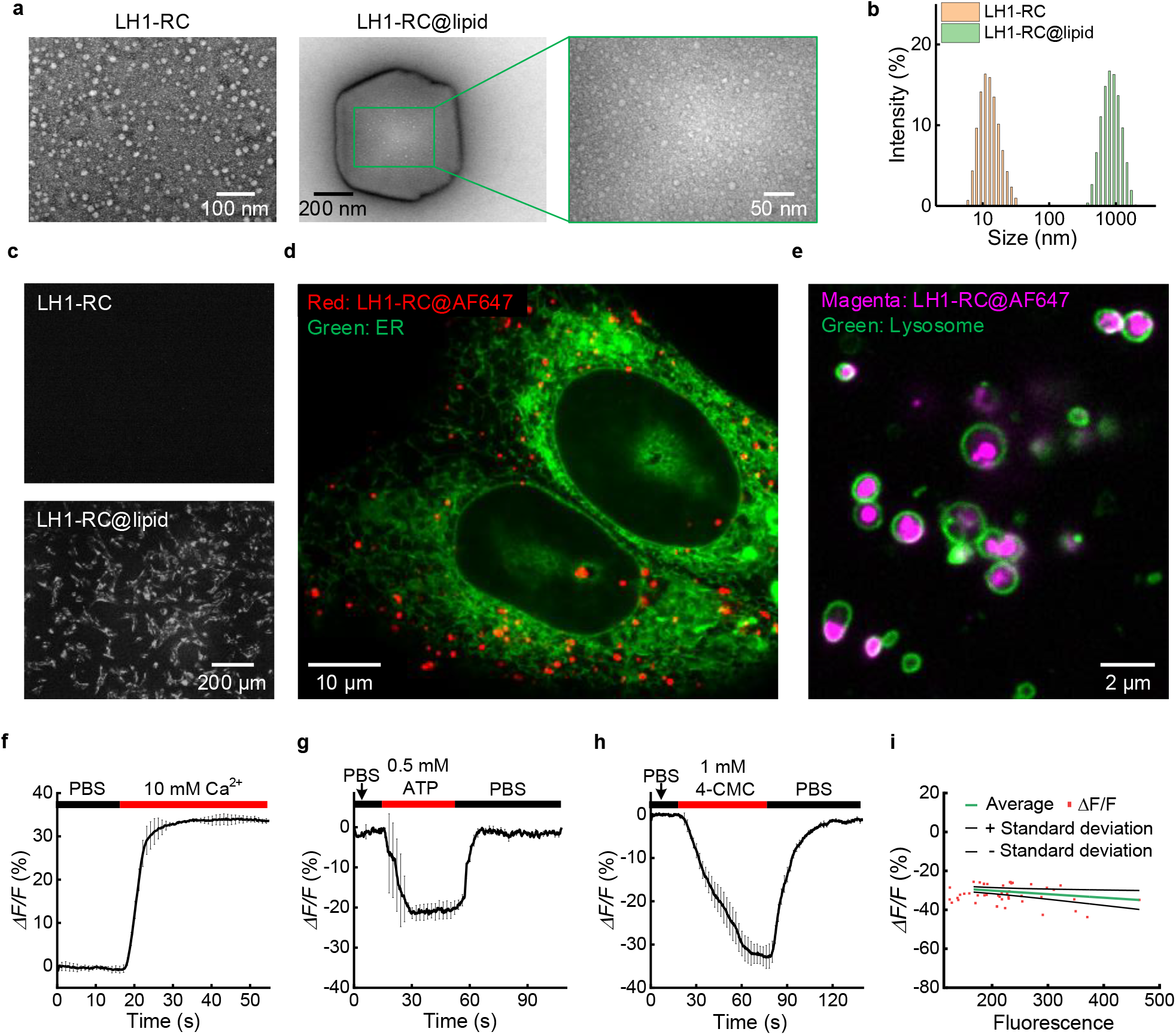
NIR-II calcium imaging under stimulation in cells. (**a**) TEM images of LH1-RC and LH1-RC@lipid. (**b**) Dynamic light scattering spectra of LH1-RC and LH1-RC@lipid in PBS. (**c**) Delivery of LH1-RC or LH1-RC@lipid into C2C12 cells at 4 hours after the addition of LH1-RC or LH1-RC@lipid to the culture medium, respectively. A 915 nm laser was used for excitation, and the NIR-II fluorescence was collected after being filtered through a 1000 nm long-pass filter. A 10× objective was used, and the exposure time was 200 ms. (**d**) Two-plex super-resolution imaging of U2-OS cells expressing VAPA-EGFP (ER, green) and incubated with LH1-RC@Alexa Fluor™ 647 (LH1-RC@AF647, red). (**e**) Two-plex imaging of U2-OS cells expressing LAMP1-mNeonGreen (lysosomes, green) and incubated with LH1-RC@AF647 (magenta). LH1-RC@AF647 was localized in lysosomes. Responses of LH1-RC in C2C12 cells to the addition of (**f**)10 mM Ca^2+^, (**g**) 0.5 mM ATP or (**h**) 1 mM of 4-CMC. *n*=3, where *n* is the number of analyzed cells. Data are shown as mean ± s.d. derived from analyzing three cells under different stimulation conditions. (**i**) Pixel by pixel plot of *ΔF/F* response from the 3 cells in Fig. 2h.

An inverted NIR-II fluorescence microscope, modified from a Nikon microscope, was used to image living cells. A 915 nm laser was used to excite the LH1-RC, and the NIR-II fluorescence was filtered with a 1000 nm long-pass filter before being collected by a camera. Fluorescence signals within the cells were observed at ∼1 hour and increased over time after the addition of LH1-RC, reaching the maximum intensity at ∼ 4 hours with a high signal-to-background ratio (Fig. 2c, Supplementary Fig.9).

To assess the localization of LH1-RC in cells, the LH1-RC was labeled with Alexa Fluor™ 647 (AF647) and delivered to U2-OS cells expressing VAPA-EGFP (an ER membrane marker, Fig. 2d) or LAMP1-mNeonGreen (a lysosome marker, Fig. 2e). U2-OS cells, a human osteosarcoma cell line, were used for organelle visualization because of their flat, spread, epithelial-like morphology and efficient exogenous gene expression^63^. Super-resolution live-cell imaging revealed that LH1-RC was internalized into lysosomes, interacting with ER, moved and trafficked together with ER tubules (Fig. 2d and Fig. 2e, Supplementary Video 1 and Supplementary Video 2). In contrast, no apparent signal was observed in cells cultured in medium containing AF647 without LH1-RC@lipid (Supplementary Fig. 10), indicating that the LH1-RC@lipid can serve as a calcium indicator for lysosomes.

The fluorescence changes of LH1-RC in response to lysosomal Ca^2+^ concentration fluctuations induced by external stimuli were characterized in the NIR-II window (Fig. 2f-i). We first added LH1-RC@lipid into the culture medium, and 6 hours later, replaced the medium with a Ca^2+^-free PBS buffer. The signal collected during the initial 10 s was used as a baseline. An apparent fluorescence signal increase (*ΔF/F* = 35.3%) was monitored after adding Ca^2+^ solution to a final concentration of 10 mM (Fig. 2f).

The lysosome is an important intracellular Ca^2+^ store with a Ca^2+^ concentration of ∼500 μM ^62^. It has been demonstrated that ATP can increase endogenous levels of nicotinic acid adenine dinucleotide phosphate (NAADP), which induces Ca^2+^ release from lysosomes^64, 65^. First, we added LH1-RC@lipid without Ca^2+^ into the culture medium. After 6 hours, the culture medium was replaced with Ca^2+^-free PBS as the imaging buffer before imaging. A decrease in peak fluorescence (– 21.7%) was observed after the addition of 0.5 mM ATP to the imaging buffer, indicating Ca^2+^ release from lysosomes. The fluorescence intensity recovered upon replacing the imaging buffer with Ca^2+^-free PBS (Fig. 2g).

Similarly, A strong decrease peak (33.0%) was observed after the addition of 1 mM 4-CMC into the imaging buffer and fluorescence intensity recovered when the imaging buffer was replaced with Ca^2+^-free PBS (Fig. 2h). Further analysis of the pixel-by-pixel plot of *ΔF/F* indicated that these cells exhibited a similar degree of release responses (Fig. 2i). This signal decrease is consistent with the 4-CMC-induced reduction of Ca^2+^ levels in the ER, as observed using a visible ER Ca^2+^ indicator^24^, and is opposite to the Fluo-4-indicated Ca^2+^ changes in the cytoplasm (Supplementary Fig. 11). Since LH1-RC was located in lysosomes and 4-CMC induces ER Ca^2+^ release through the agonist ryanodine receptor (RyR)^66^, we inferred that the Ca^2+^ release from lysosomes induced by 4-CMC arises from the crosstalk between Ca^2+^ release from the lysosome and the ER at membrane contact sites^65^.

### NIR-I and NIR-II phantom imaging of calcium-induced fluorescence changes

Phantom imaging of the near-infrared calcium indicators iGECI^22^ and LH1-RC was performed in NIR-I and NIR-II windows (Fig. 3). A pair of 300-μm-diameter capillaries, separated by ∼ 300 μm and filled with iGECI or LH1-RC with or without Ca^2+^ (Fig. 3a), were imaged through a 1% intralipid solution layer of varying thickness to mimic muscle tissues^43^. NIR-I fluorescence of iGECI was collected after being filtered by a 700 nm long-pass (LP) filter, while NIR-II fluorescence of LH1-RC was filtered using long-pass filters with cut-on wavelengths ranging from 1000 to 1300 nm (Fig. 3b and Supplementary Fig. 12). The fluorescence of iGECI decreases in the presence of Ca^2+^, while it increases when Ca^2+^ is removed, due to FRET between the miRFP670 donor and the miRFP720 acceptor^22^. For NIR-I imaging, two iGECI-filled capillaries, with or without Ca^2+^, were individually resolved through an intralipid layer of up to ∼ 3 mm (Fig. 3b, first column; Supplementary Fig. 12, first column). The *ΔF/F* value quickly decreased from -20.5% to nearly zero as the intralipid depth increased from 0 to 3 mm (Fig. 3c). Moreover, as the thickness of the intralipid layer exceeded 3 mm, the capillary signal gradually approached the background level (Fig. 3e), indicating the maximum penetration depth of the iGECI signal is ∼ 3 mm.

**Figure 3.**
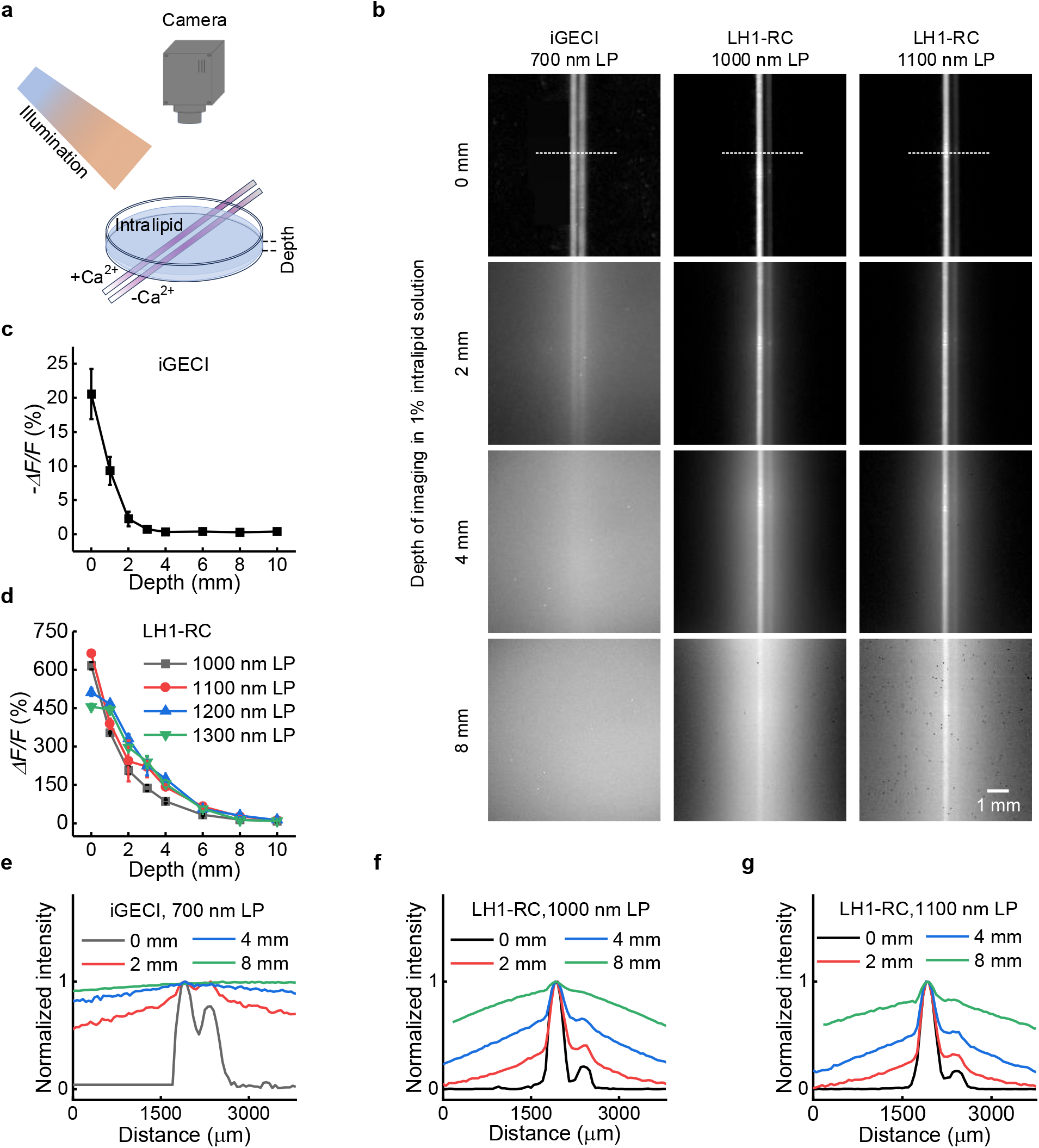
Phantom imaging in NIR-I and NIR-II sub-regions. (**a**) Schematic of the experimental setup for phantom imaging of two capillaries. (**b**) Fluorescence imaging of 300-μm-diameter capillary tubes filled with 1.8 µM of iGECI or LH1-RC, with or without Ca^2+^, immersed at different depths in a 1% intralipid solution using a wide-field imaging system. For iGECI imaging, a 660 nm laser was used for excitation, and a 700 nm long-pass filter was applied to filter the fluorescence signal. For LH1-RC imaging, a 915 nm laser was used for excitation, and the NIR-II fluorescence was filtered using long-pass filters with cut-on wavelengths of 1000 nm and 1100 nm. (**c**) Calcium-induced fluorescence changes of iGECI imaged through 1% intralipid solutions with different thicknesses in the NIR-I window. *ΔF/F= (F*_+Ca_ *-F*_-Ca_*)/F*_-Ca_, where *F*_+Ca_ and *F*_-Ca_ represent fluorescence signals of iGECI with or without Ca^2+^, respectively. (**d**) The fluorescence changes of LH1-RC imaged through 1% intralipid solutions in different NIR-II sub-windows. *ΔF/F= (F*_+Ca_ *-F*_-Ca_*)/F*_-Ca_. Normalized fluorescence intensity profiles of (**e**) iGECI-filled or (**f,g**) LH1-RC-filled capillaries at different intralipid depths along the dotted line shown in Fig. 3b. (**c,d**) Error bars represent s.d. for *n* = 3.

Two capillaries filled with LH1-RC, with or without Ca^2+^, were individually observed in the 1000-1700 nm NIR-II window even beyond 8 mm, with a *ΔF/F* of 615.9% without intralipid and a *ΔF/F* of 14.6% through an 8-mm intralipid layer (Fig. 3b, second column; Fig. 3d; Supplementary Fig. 12, second column). NIR-II imaging using a 1200 nm long-pass filter allowed better contrast and achieved a higher *ΔF/F* of 30.1% at an immersion depth of 8 mm (Fig. 3b, fourth column; Fig.3d; Supplementary Fig. 12, fourth column). *ΔF/F* at an 8 mm penetration depth can be further increased by using longer filters, while this comes at the expense of the fluorescence intensity (Fig. 3d and Supplementary Fig. 12). NIR-II imaging offers better contrast, resolution, and penetration depth than NIR-I imaging. LH1-RC provides greater fluorescence changes, ∼ 3-4 times deeper penetration (Supplementary Fig. 12), and better signal-to-noise ratios (Supplementary Fig. 13) compared to previous iGECI^22^. Therefore, NIR-II imaging of LH1-RC facilitates the detection of Ca^2+^ signaling in deep tissues.

### *In vivo* NIR-II calcium imaging of intact human tumors

Non-invasive *in vivo* calcium imaging of intact tumors in their native physiological environment can reflect the actual state of cellular activities and is promising but challenging due to current technological limitations^67^. NIR-II calcium imaging using LH1-RC offers a potential approach for *in vivo* tumor investigation, facilitated by its deeper penetration, improved resolution, and high sensitivity to Ca^2+^ changes.

We first performed NIR-II calcium imaging of intact HeLa human cervical tumors using LH1-RC with a wide-field imaging system reflecting global responses (Fig. 4b) and a light-sheet microscope (LSM) ^39, 42^ for the observation of individual cell calcium signaling (Fig. 4e). Eight hours post intratumoral (i.t.) injection of LH1-RC@lipid (Fig. 4a), strong fluorescence signals were collected after being filtered by a 1000 nm long-pass filter under 915 nm excitation (Fig. 4c, f). No apparent spontaneous calcium transients were observed by the wide-field imaging system (Supplementary Video 3) and LSM (Supplementary Video 4), consistent with previous *in vitro* imaging results^67^.

**Figure 4.**
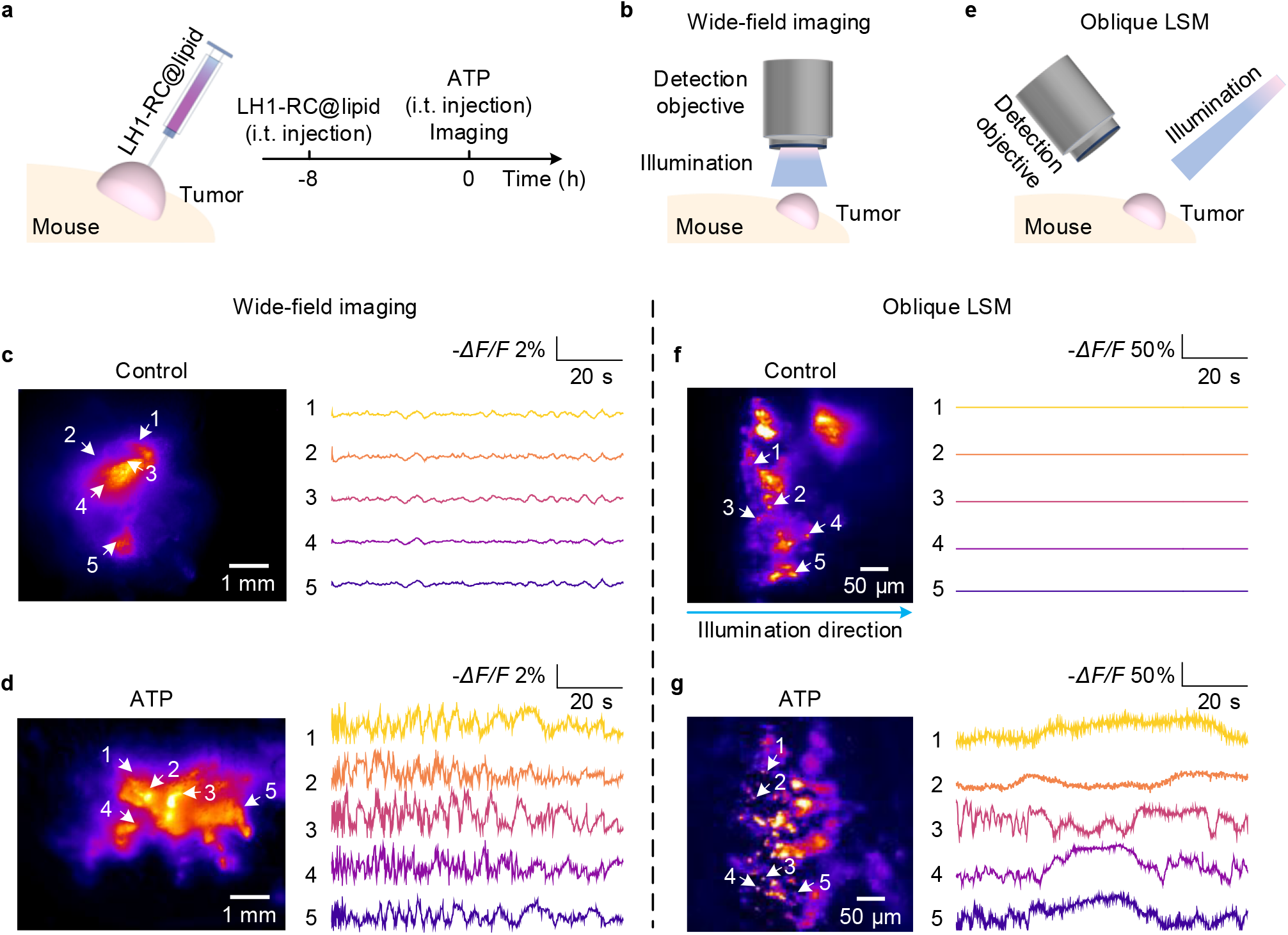
Non-invasive NIR-II calcium imaging of HeLa tumors. (**a**) Intratumoral injection of LH1-RC@lipid and imaging schedule. For the control group, LH1-RC@lipid was injected intratumorally (i.t.) ∼8 hours before imaging. For the treatment group, NIR-II calcium imaging was performed immediately post the i.t. injection of ATP and ∼ 8 hours after the i.t. injection of LH1-RC@lipid. (**b**) A simplified schematic of the NIR-II wide-field imaging system. (**c**) Wide-field calcium imaging of a HeLa tumor in the control group of mice that received no drug treatment. Similar results were observed in *n* = 3 mice. (**d**) Wide-field calcium imaging of a HeLa tumor under 1 µM ATP stimulation. For the wide-field imaging, a 915 nm laser was used for excitation, and fluorescence was collected after being filtered by a 1000 nm long-pass filter. The exposure time was 100 ms. Similar results were observed in *n* = 3 mice. (**e**) A simplified schematic of NIR-II LSM. NIR-II LSM of calcium signals in the HeLa tumors labeled by LH1-RC@lipid in (**f**) the control group of mice with no treatment, and (**g**) the group of mice treated with 1 µM ATP. Similar results were observed in *n* = 2 mice for the control group and the treatment group. The light sheet was positioned at the location of the tumor protrusion in the mouse. A 915 nm laser was used for excitation, and fluorescence was collected in the 1000-1700 nm window. A 5× objective and a 10× objective were used for excitation and detection, respectively. The exposure time was100 ms. (**c,d,f,g**) Regions of interest (ROIs) were selected randomly for analysis.

To evaluate the capability of LH1-RC for NIR-II calcium imaging, 1 µM ATP was intratumorally injected 8 hours post injection of LH1-RC@lipid (Fig. 4a). Wide-field imaging shows a series of fluorescence changes with a highest -*ΔF/F* of 3.6%, which is ten times higher than the previous NIR-I imaging of NIR-GECO1^21^ (Fig. 4d, Supplementary Fig. 14a). Different areas exhibited varying fluctuation patterns, generating Ca^2+^ waves within the tumor (Supplementary Video 5). This was further confirmed by LSM-based calcium imaging at the cellular resolution (Fig. 4g, Supplementary Video 6, Supplementary Fig. 14b). LSM calcium imaging achieved a much larger *-ΔF/F* of 82.1% than wide-field microscopy, benefiting from the suppressed background in LSM.

### *In vivo* NIR-II calcium imaging during tumor treatment

Dexamethasone (Dex) regulates TRPV6 (Transient Receptor Potential channel family, Vanilloid subfamily member 6) calcium channel expression, increases intracellular calcium concentration^68^, and shows anticancer efficacy in lung cancers clinically^69^. In contrast, SOR-C13 is a high-affinity TRPV6 antagonist that inhibits tumor growth by targeting the TRPV6 calcium channel and is undergoing clinical trials^70^. To evaluate the influence of Dex and SOR-C13 on calcium signaling in tumors, we performed *in vivo* NIR-II calcium imaging of A549 tumors over the course of treatment. A549 cells are human non-small cell lung cancer cells and exhibit spontaneous intracellular calcium transients^45, 46^. A549 tumor was inoculated on the hindlimb of BALB/c nude mice (Fig. 5a). LH1-RC@lipid was utilized to enhance LH1-RC delivery efficiency to A549 cells (Supplementary Figs. 15-17). Both the groups treated and not treated with LH1-RC@lipid showed similar cell viability (> 90%), demonstrating negligible cytotoxicity of LH1-RC@lipid to A549 cells (Supplementary Fig. 18). LH1-RC@lipid was injected intratumorally 8 hours prior to the treatment with Dex or SOR-C13 (Fig. 5a, b). Inflammatory cytokine levels of IL-6 and TNF-α in A549 tumor-bearing mice were assessed after injecting 50 µL of PBS or 0.03 µg/µL LH1-RC@lipid at different time points (0 h, 24 h, and 48 h). Mild inflammation was induced by injecting LH1-RC@lipid at 24 hours, which decreased to a level slightly higher than that of the PBS-injected group at 48 hours post injection. No significant differences were observed among the groups (based on Pair-sample T-test, *n* = 3), indicating that LH1-RC@lipid has a low immunogenicity risk (Supplementary Fig. 19).

**Figure 5.**
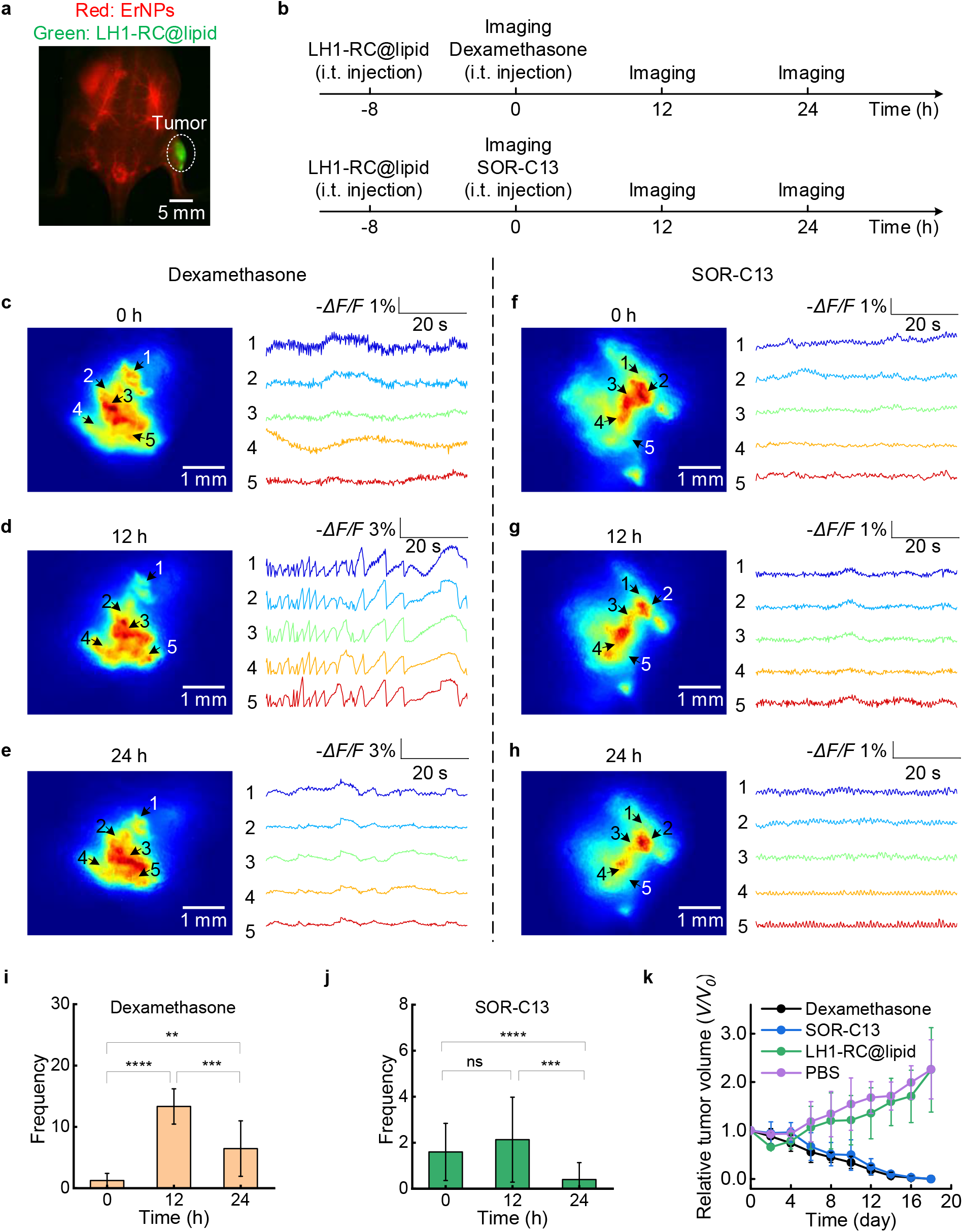
Non-invasive NIR-II calcium imaging of A549 tumors in response to different treatment strategies. (**a**) Wide-field imaging of A549 tumor labeled with LH1-RC@lipid (green) and ErNPs filling vessels (red) 24 hours post intratumoral injection of LH1-RC@lipid and 5 min post intravenous injection of ErNPs. (**b**) Treatment and imaging schedule. The first NIR-II calcium imaging of A549 tumors was performed 8 hours after the i.t. injection of LH1-RC@lipid. Following the initial imaging at 0 h, A549 tumors were treated with 10 nM dexamethasone or 640 µM SOR-C13. The calcium transients in response to these treatments were then monitored at 12 h and 24 h. Wide-field calcium imaging of A549 tumors (**c**) before dexamethasone treatment at 0 h, and after 10 nM dexamethasone treatment at (**d**) 12 h and (**e**) 24 h. NIR-II wide-field calcium imaging of A549 tumors (**f**) before SOR-C13 treatment, and after 640 µM SOR-C13 treatment at (**g**) 12 h and (**h**) 24 h. A 915 nm laser was used for excitation, and fluorescence was collected in the 1000-1700 nm window. The exposure time was 200 ms. Calcium signal frequency of A549 tumors in response to the treatment with (**i**) 10 nM dexamethasone and (**j**) 640 µM SOR-C13 at different time points. Similar results were observed in *n* = 3 mice for each treatment. (**k**) The relative tumor volume (*V/V*_*0*_, where *V*_*0*_ is the tumor volume on Day 0, *V* is the tumor volume) of A549 tumor-bearing mice was measured after different treatments. Similar results were observed in *n* = 3-6 mice for each group. Day 0 is defined as 4 days after tumor inoculation, with the tumor size at Day 0 being ∼100 mm^3^. (**c-e,f-h**) ROIs were selected randomly for analysis. (**i,j**) *P < 0.05, **P < 0.01, ***P < 0.001, ****P < 0.0001 as measured by Pair-sample T-test. ns indicates not significant.

In the Dex treatment group, A549 tumors exhibited slow spontaneous calcium transients with a highest -*ΔF/F* of 1.5% before treatment (Fig. 5c; Supplementary Video 7). Higher-frequency calcium signals were observed at 12 and 24 hours post-intratumoral injection of Dex (Fig. 5d,e,i; Supplementary Videos 8 and 9). A larger -*ΔF/F* of 4.8% and 2.6% was monitored 12 hours and 24 hours after Dex injection, respectively (Fig. 5d,e). In the SOR-C13 treatment group, A549 tumors exhibited slow spontaneous calcium transients before treatment, similar to those observed in the Dex group (Fig. 5c,f; Supplementary Video 10). We observed a slight increase in calcium transient frequency 12 hours post SOR-C13 injection, but statistical analysis showed no significant difference compared to that at 0 hours. The calcium transients were suppressed 24 hours after SOR-C13 injection (Fig. 5g,h,j; Supplementary Videos 11 and 12). Compared to the control group with tumors treated with PBS or LH1-RC@lipid, tumors in the Dex and SOR-C13 treatment groups showed noticeable shrinkage (Fig. 5k, Supplementary Figs. 20 and 21). Although Dex and SOR-C13 induced different changes in calcium signaling within A549 tumors, both could contribute to the tumor treatment process.

## Conclusion

Non-invasive longitudinal NIR-II calcium imaging of intact tumors with high sensitivity, resolution, and contrast has been demonstrated. The naturally derived LH1-RC from *Tch. tepidum* contains 16 calcium-binding sites and 32 Bchl *a* molecules, affording high sensitivity to Ca^2+^ and strong NIR-II emission in the NIR-II range of 1000-1300 nm. The pronounced red-shifted absorption of LH1-RC from *Tch. tepidum* is primarily attributed to its coordination of 16 calcium ions, distinguishing it from other LH1-RC complexes^55, 57, 71^. This Ca^2^+ binding not only enhances spectral red-shifting but also imparts exceptional thermal stability. The conformation of LH1-RC provides an alternative mechanism for Ca^2+^ sensing, distinct from traditional calcium-indicating proteins that rely on calmodulin (CaM)^21, 22^. The photophysical properties of LH1-RC, including charge separation and electron transport in the RC complex, are unlikely to exert unintended effects on host cell physiology, as the purified LH1-RC cannot complete the full electron transfer process in the mammalian host environment^72^. LH1-RC exhibits excellent stability and is suitable for one-photon NIR-II wide-field imaging as well as light-sheet microscopy, providing a complementary method for deep-tissue calcium imaging using two-/multi-photon microscopy in combination with visible and NIR-I calcium indicators. Compared with two-or multi-photon imaging using visible-wavelength Ca^2+^ indicators, LH1-RC-based one-photon NIR-II imaging enables multiscale imaging, from cellular resolution to a large field of view, offering deep penetration and fast imaging speed. LH1-RC also allows non-invasive in vivo imaging without the need for surgery or implanting a cranial window in mice (Supplementary Table 3). While LH1-RC was validated in subcutaneous tumors for calcium imaging, it is also readily applicable to deeper or metastatic tumors. LH1-RC emits bright fluorescence in the NIR-II window, facilitating deep tissue penetration. Ligand-mediated targeting could be applied to enhance the specificity of LH1-RC for deep tumors or metastases in the future.

Currently, LH1-RC is not genetically encodable, limiting its use to exogenous delivery scenarios. Targeted delivery strategies can be designed to enhance the transport of LH1-RC to specific cells. It is also attractive to pursue molecular engineering strategies to convert LH1-RC into a genetically encoded calcium indicator and optimize its gene sequence to be compatible with the genomes of mammalian cells. The structure of LH1-RC can be further optimized to enhance its biocompatibility with mammalian cells and its sensitivity to Ca^2^+. For example, the reaction center of the LH1-RC complex can be genetically knocked out to disrupt the photosynthetic process^56^, allowing LH1 to function solely as a light-harvesting antenna and avoiding the potential impact of charge separation-generated electrons on the host organism. This modification has the potential to enable LH1 to serve as an excellent NIR-II calcium indicator for *in vivo* imaging. NIR-II calcium imaging can be further enhanced by developing brighter fluorescent proteins and synthetic fluorophores with longer excitation and emission wavelengths. NIR-II calcium imaging paves the way for advanced research in both neuroscience and oncology.

## Data availability

The data that support the findings of this study are available from the corresponding author upon request.

## Methods

### LH1-RC preparation

*Tch. tepidum* were cultured anaerobically at 48 °C for 7 days. The harvested bacteria were disrupted by sonication in a 20 mM Tris-HCl buffer at a pH of 8.5. Chromatophores were initially treated with 0.35% (w/v) lauryldimethylamine N-oxide at 25 °C for 60 min, followed by centrifugation at 150,000 × *g* for 90 min. The pellet was subsequently treated with 1.0% (w/v) *n*-Octyl-β-D-Glucopyranoside (OG) and ultracentrifuged at 150,000 × *g* for 90 min to extract the LH1-RC. The supernatant was loaded onto a DEAE anion-exchange column (Toyopearl 650S, Tosoh Corp.) equilibrated at 4 °C with 20 mM Tris-HCl buffer, pH 7.5, containing 0.8% (w/v) OG. The LH1-RC fraction was eluted by a linear gradient of CaCl_2_ from 0 to 50 mM, and peak fractions with an *A*_915_/*A*_280_ ratio over 2.10 were collected^52^. *A*_915_ and *A*_280_ are the absorbance values of the sample at 915 nm and 280 nm, representing the absorbance of LR proteins and total proteins in the sample, respectively.

### Preparation of iGECI

iGECI and biliverdin were expressed in the BL21-AI host (Beyotime). Bacteria were grown in Lysogeny Broth (LB) medium supplemented with ampicillin for 6-8 h, followed by induction of protein expression with 0.1% arabinose. The iGECI proteins were purified using Ni-NTA agarose (Beyotime).

### Absorbance measurement

A microplate reader (Thermo Fisher) was used to measure the absorbance of LH1-RC and iGECI. To characterize the response of LH1-RC to different Ca^2+^ concentrations, a 199 µL solution containing 1.2 µM LH1-RC was added to a 96-well plate. Then, 1 µL of Ca^2+^ solution at different concentrations (4 mM, 8 mM, 12 mM, 14 mM, 20 mM, 40 mM, 200 mM) was dropped into the LH1-RC-containing solution to achieve different Ca^2+^ concentrations (20 µM, 40 µM, 60 µM, 70 µM, 100 µM, 200 µM, 1000 µM). Absorbance intensities were plotted against Ca^2+^ concentrations and fitted by a sigmoidal binding function to determine the Hill coefficient and *K*_d_^21^.

### Quantum yield measurement

The quantum yield of LH1-RC with 1 mM Ca^2+^ in 20 mM Tris-HCl buffer at a pH of 7.5 was measured using a fluorescence spectrometer (Edinburgh, FLS1000) equipped with an integrating sphere. The excitation wavelength was 915 nm, and the signal acquisition range was from 1000 nm to 1400 nm.

### NIR-I and NIR-II phantom imaging

A pair of 300-µm diameter capillaries was immersed in 1% intralipid at varying depths (0-10 mm). To image the iGECI-containing capillaries, a 660 nm laser was used for excitation. The fluorescence was filtered by a 700 nm long-pass filter and recorded by an Andor visible camera. To image capillaries filled with LH1-RC, a 915 nm laser was applied for excitation. Fluorescence was filtered by 1000 nm, 1100 nm, 1200 nm, or 1300 nm long-pass filters before being recorded by an indium gallium arsenide (InGaAs) camera.

### Lipid encapsulation strategy of LH1-RC for cultured C2C12 cells

1.5 µL of Lipofectamine™ 3000 reagent (Thermo Fisher) was diluted in 25 µL of Opti-MEM Medium (tube 1), and 1 µg of LH1-RC without Ca^2+^ was diluted in 25 µL of Opti-MEM (tube 2). Tube 2 was then added to tube 1. The mixture was incubated for 10-15 minutes at room temperature.

### LH1-RC@lipid preparation for TEM

100 µL of 0.03 µg/µL LH1-RC@lipid in PBS buffer was prepared. The samples were identified by negative staining and observed under transmission electron microscopy (TEM). First, a 200-mesh copper grid (Quantifoil® R 2/2) was placed in a glow emission cleaning instrument (PELCO easiGlow), where a 60-second vacuum procedure was followed by a 30-second glow discharge at a power of 60 W. Then, 5 μL of the sample was dripped onto the cleaned grid and allowed to stand for 60 seconds to ensure the sample fully combined with the grid. Use filter papers to gently remove excess unbound samples. Subsequently, 5 μL of uranium acetate (2%) was added to the grid allowed to stand for 60 seconds. Excess dye was gently removed with filter paper again. The stained grid was transferred to a sample box for storage and subsequent observation.

### LH1-RC delivery and NIR-II calcium imaging *in vitro*

C2C12 cells were cultured in growth medium consisting of DMEM supplemented with 10% fetal bovine serum (FBS) and antibiotics. When C2C12 cells reached 70-80% confluency, they were passaged by detaching them with trypsin-EDTA, neutralizing the trypsin, and then seeding them into a 24-well plate. After 16 hours of culture, LH1-RC@lipid was added to each well. Fluorescence images were recorded at different time points by an inverted NIR-II fluorescence microscope modified from a Nikon microscope. A 10× objective was used. Before NIR-II imaging, cells were washed three times with PBS buffer without Ca^2+^.

Alexa Fluor™ 647 Conjugation Kit (abcam, ab269823) was used to label LH1-RC^73^. First, 100 µg LH1-RC protein was diluted into 100 µL PBS. 1-2 µL of modifier was added to the protein solution and mixed thoroughly. The resulting mixture was then added to AF647 dye and incubated for 15 minutes at room temperature with gentle mixing to facilitate conjugation. Following the conjugation reaction, the reaction was quenched by the addition of 1-2 µL of quencher, followed by a further 5-minute incubation. The labeled protein was used immediately for subsequent experiments.

U2-OS cells transfected with pEGFPC1-hVAP-A (Addgene,104447) or LAMP1-mNeonGreen (Addgene, 98882) were treated with LH1-RC@AF647 and imaged on a VT-iSIM super-resolution imaging system (Visitech International), based on an Olympus IX83 microscope, with a 100×/1.45 objective (UPLXAPO100XO). Cells were imaged in a stage incubator, maintained at 37 °C and 5% CO_2_. Simultaneous imaging of pEGFPC1-hVAP-A/LAMP1-mNeonGreen and LH1-RC@AF647 was performed using 488 nm laser excitation with an ET525/50 nm emission filter (Chroma) and 640 nm laser excitation with an ET700/75 nm emission filter (Chroma), respectively.

For NIR-II calcium imaging of C2C12 cells, the cells were washed three times with Ca^2+^-free PBS buffer, which was also used as the imaging buffer. Cells were illuminated with a 915 nm laser. A 10× objective was used. Fluorescence was filtered by a 1000 nm long-pass filter and recorded by an InGaAs camera while C2C12 cells were stimulated by different drugs (10 mM CaCl_2_, 0.5 mM ATP or 1 mM 4-CMC).

### Lipid encapsulation strategy of LH1-RC for cultured C2C12 cells or NIR-II imaging *in vivo*

3 µg of LH1-RC and 5 µL of Cas9 (Thermo Fisher) were diluted in 50 µL of Opti-MEM medium (tube 1). 3 µL of CRISPRMAX™ (Thermo Fisher) was diluted in 50 µL of Opti-MEM (tube 2). Then tube 1 was added to tube 2. The mixture was incubated for 5-10 minutes at room temperature. Lipofectamine™ CRISPRMAX™ is another transfection reagent that demonstrates superior delivery efficiency^74^; therefore, we used it for the *in vivo* delivery of LH1-RC to A549 cells.

### Mouse handling, tumor inoculation, and treatment

All animal experiments were approved by the University of Hong Kong’s Committee on the Use of Live Animals in Teaching and Research (CULATR). BALB/c nude female mice were purchased from the Centre for Comparative Medicine Research, the University of Hong Kong. The ambient relative humidity was maintained at 55-65%, and the temperature was ∼25 °C. Before NIR-II *in vivo* imaging, a rodent anesthesia machine with 3 L/min O_2_ gas flow mixed with 2% isoflurane was used to anesthetize the mice. During imaging, the mouse was kept anesthetized by a nose cone delivering 1.5 L/min O_2_ gas mixed with 0.3% isoflurane.

Mice were randomly selected from cages for all experiments. Each group in the study consisted of 3 to 6 mice. HeLa tumors were induced by subcutaneous (s.c.) injection of 1 × 10^7^ cells suspended in 100 μL PBS. A549 tumors were established by s.c. injection of 1 × 10^7^ cells suspended in 100 μL PBS.

For *in vivo* imaging of HeLa tumors, the experiment was performed 8 days after tumor inoculation, when the tumors reached ∼200 mm^3^ in volume. 50 µL of 0.03 µg/µL LH1-RC@lipid was injected intratumorally 8 hours before imaging to ensure delivery of the LH1-RC@lipid into the tumor cells. 20 µL of 1 µM ATP was injected into the HeLa tumors before imaging. Control group mice received no drug treatment and were imaged 8 hours after LH1-RC@lipid delivery.

For *in vivo* imaging of A549 tumors, 50 µL of 0.03 µg/µL LH1-RC@lipid was injected intratumorally 8 hours before imaging to ensure delivery of the LH1-RC@lipid into the tumor cells. The tumors were imaged before any treatment to obtain baseline calcium signals in A549 tumors at 0 hour. After imaging at 0 h, mice were intratumorally injected with 20 µL of 10 nM dexamethasone or 640 µM SOR-C13 for the treatment groups. They were subsequently imaged at 12 h and 24 h, respectively.

For the treatment of A549 tumors, the experiment was performed 4 days after tumor inoculation, when the tumors reached ∼100 mm^3^ in volume. The mice were randomly divided into four groups and intratumorally injected with dexamethasone, SOR-C13, LH1-RC@lipid, or PBS. Tumor size was recorded every 2 days. Tumor volume was calculated using the formula *V=*0.52*×a×b*^*2*^, where *a* is the maximum length of the tumor, *b* is the maximum width of the tumor.

### NIR-II wide-field imaging *in vivo*

*In vivo* imaging of tumor-bearing mice as shown in Fig. 5a was performed using a 2D InGaAs camera. For the ErNPs^45^ channel, excitation light was provided by a 980 nm laser, and fluorescence was collected after being filtered by a 1500 nm long-pass filter. For the LH1-RC channel, a 915 nm laser was used for excitation, and fluorescence was collected after being filtered by a 1000 nm long-pass filter. The actual excitation intensity of the 980 nm and 915 nm lasers at the imaging plane was 26.3 mW cm^-2^ and 43.1 mW cm^-2^, respectively. The exposure times for imaging were 50 ms for ErNPs and 30 ms for LH1-RC.

Tumors shown in Fig. 4c, d and Fig. 5c-h were imaged using a combination of a 150-mm achromatic lens and a 200-mm tube lens. A 915 nm laser with an actual excitation intensity of 45.5 mW cm^-2^ was used for excitation. Fluorescence was collected after being filtered by a 1000 nm long-pass filter. The exposure time for the InGaAs camera was 100-200 ms.

After intratumoral injection and sectioning of the injected tumor, we observed LH1-RC@lipid signals in each tumor slice, but the signals were not evenly distributed, which could be attributed to tumor heterogeneity (Supplementary Fig. 6).

### Light-sheet microscopy *in vivo*

For LSM imaging, the illumination and detection objectives are perpendicular to each other and aligned at a 45° angle to the sample^42^. The light sheet was positioned at the location of the tumor protrusion in the mouse. Fluorescence was collected using an InGaAs camera after being filtered by a 1000 nm long-pass filter. We used a 5× objective for excitation and a 10× objective for imaging. A 915 nm laser filtered by a 905-925 nm bandpass filter was used for excitation. The exposure time was 100-200 ms.

## Data processing

The raw image data recorded by an InGaAs camera was processed using ImageJ (2.14.0). Multicolor fluorescence images were overlaid with ImageJ. The standard deviation and mean were calculated by Origin 2021. *ΔF/F=(F−F*_baseline_*)/F*_baseline_, where *F* is the real-time fluorescence intensity during the imaging period. For cell imaging, *F*_baseline_ is the average intensity during the first 10 s. For *in vivo* imaging, *F*_baseline_ was created by the Create Baseline function in Origin 2021, as described in reference^75^ . All curves were corrected for bleaching using Origin 2021. A MATLAB program was used to align fluorescence images captured by a visible camera and an InGaAs camera.

## Acknowledgments

This study was supported by the JC STEM Lab of Nanoscience and Nanomedicine and the JC STEM Lab of Molecular Imaging funded by The Hong Kong Jockey Club Charities Trust, General Research Fund (RGC No. 17212424, 17102722, 17300523, 17302724), Early Career Scheme (RGC No. 27204623) from the Research Grants Council of Hong Kong SAR, and National Natural Science Foundation of China (Project No. T2522030).

## Author contributions

F.W., H.D. and D.X. conceived and designed the experiments. L.Y., X.Y. and G.W. provided purified LH1-RC. D.X. performed the experiments. Z.W. set up the wide-field imaging system. H. Y., G. C. and H. J. labeled the LH1-RC and performed the U2-OS cell imaging. D.X., Z.W., S.X., H.Y., Y.L., Z.D. L.Y. H.D. and F.W analyzed the data. D.X., L.Y. H.D. and F.W. wrote the manuscript. All authors contributed to the general discussion and revision of the manuscript.

## Materials & Correspondence

Correspondence and requests for materials should be addressed to F.W.

