## Supplementary Materials for "*In Vivo* Calcium Imaging in the Near-Infrared II Window"

**SUPPLEMENTARY VIDEO CAPTIONS**

**Supplementary Video 1 |** Time-course two-plex imaging of LH1-RC@AF647 (red) and ER (green). Simultaneous imaging of ER and LH1-RC@AF647 was performed using 488 nm laser excitation with an ET525/50 nm emission filter, and 640 nm laser excitation with an ET700/75 nm emission filter, respectively. A VT-iSIM super-resolution imaging system, based on an Olympus IX83 microscope equipped with a 100×/1.45 objective was used.

**Supplementary Video 2 |** Time-course two-plex imaging of LH1-RC@AF647 (magenta) and lysosomes (green). Simultaneous imaging of lysosomes and LH1-RC@AF647 was conducted using 488 nm laser excitation with an ET525/50 nm emission filter, and 640 nm laser excitation with an ET700/75 nm emission filter, respectively. A VT-iSIM super-resolution imaging system, based on an Olympus IX83 microscope equipped with a 100×/1.45 objective was used.

**Supplementary Video 3 |** Non-invasive, time-course NIR-II calcium imaging of a HeLa tumor was performed using a wide-field imaging system 8 hours after intratumoral injection of LH1-RC@lipid (as shown in Fig. 4c). A 915 nm laser was used for excitation and fluorescence was collected after being filtered by a 1000 nm long-pass filter. The exposure time was 100 ms. No apparent spontaneous calcium transients were observed.

**Supplementary Video 4 |** Time-course NIR-II wide-field calcium imaging of a HeLa tumor treated with 1 μM ATP (as shown in Fig. 4d). LH1-RC@lipid was injected intratumorally 8 hours before the intratumoral administration of ATP and wide-field imaging. A 91 5 nm laser was used for excitation, and fluorescence was collected after being filtered by a 1000 nm long-pass filter. The exposure time was 100 ms. Fast and intensive calcium transients were observed.

**Supplementary Video 5 |** Non-invasive time-course NIR-II calcium imaging of a HeLa tumor was performed using LSM 8 hours after intratumoral injection of LH1-RC@lipid (as shown in Fig. 4f). A 915 nm laser, filtered by a 905-925 nm bandpass filter, was used for excitation. Fluorescence was collected using an InGaAs camera after being filtered by a 1000 nm long-pass filter. We used a 5× objective for excitation and a 10× objective for imaging. The exposure time was 100 ms. No apparent spontaneous calcium transients were observed by the NIR-II LSM.

**Supplementary Video 6 |** Instant non-invasive time-course NIR-II calcium imaging of a HeLa tumor treated with 1 μM ATP was performed using LSM 8 hours after intratumoral injection of LH1-RC@lipid (as shown in Fig. 4g). A 915 nm laser, filtered by a 905-925 nm bandpass filter, was used for excitation. Fluorescence was collected using an InGaAs camera after being filtered by a 1000 nm long-pass filter. We used a 5× objective for excitation and a 10× objective for imaging. The exposure time was 100 ms. Calcium imaging at cellular resolution was achieved by the NIR LSM.

**Supplementary Video 7 |** Non-invasive time-course calcium imaging of an A549 tumor was conducted using a wide-field imaging system 8 hours after intratumoral injection of LH1-RC@lipid before dexamethasone treatment (as shown in Fig. 5c). A 915 nm laser was used for excitation and fluorescence was collected after being filtered by a 1000 nm long-pass filter. The exposure time was 200 ms.

**Supplementary Video 8 |** Non-invasive time-course calcium imaging of an A549 tumor was performed by a wide-field imaging system 20 hours after intratumoral injection of LH1-RC@lipid and 12 hours after intratumoral administration of dexamethasone (as shown in Fig. 5d). A 915 nm laser was used for excitation and fluorescence was collected after being filtered by a 1000 nm long-pass filter. The exposure time was 200 ms.

**Supplementary Video 9 |** Non-invasive time-course calcium imaging of an A549 tumor was performed by a wide-field imaging system 32 hours after intratumoral injection of LH1-RC@lipid and 24 hours after intratumoral administration of dexamethasone (as shown in Fig. 5e). A 915 nm laser was used for excitation and fluorescence was collected after being filtered by a 1000 nm long-pass filter. The exposure time was 200 ms.

**Supplementary Video 10 |** Non-invasive time-course calcium imaging of an A549 tumor was conducted using a wide-field imaging system 8 hours after intratumoral injection of LH1-RC@lipid before SOR-C13 treatment (as shown in Fig. 5f). A 915 nm laser was used for excitation and fluorescence was collected after being filtered by a 1000 nm long-pass filter. The exposure time was 200 ms.

**Supplementary Video 11 |** Non-invasive time-course calcium imaging of an A549 tumor was performed by a wide-field imaging system 20 hours after intratumoral injection of LH1-RC@lipid and 12 hours after intratumoral administration of SOR-C13 (as shown in Fig. 5g). A 915 nm laser was used for excitation and fluorescence was collected after being filtered by a 1000 nm long-pass filter. The exposure time was 200 ms.

**Supplementary Video 12 |** Non-invasive time-course calcium imaging of an A549 tumor was performed by a wide-field imaging system 32 hours after intratumoral injection of LH1-RC@lipid and 24 hours after intratumoral administration of SOR-C13 (as shown in Fig. 5h). A 915 nm laser was used for excitation and fluorescence was collected after being filtered by a 1000 nm long-pass filter. The exposure time was 200 ms.

### SUPPLEMENTARY FIGURES

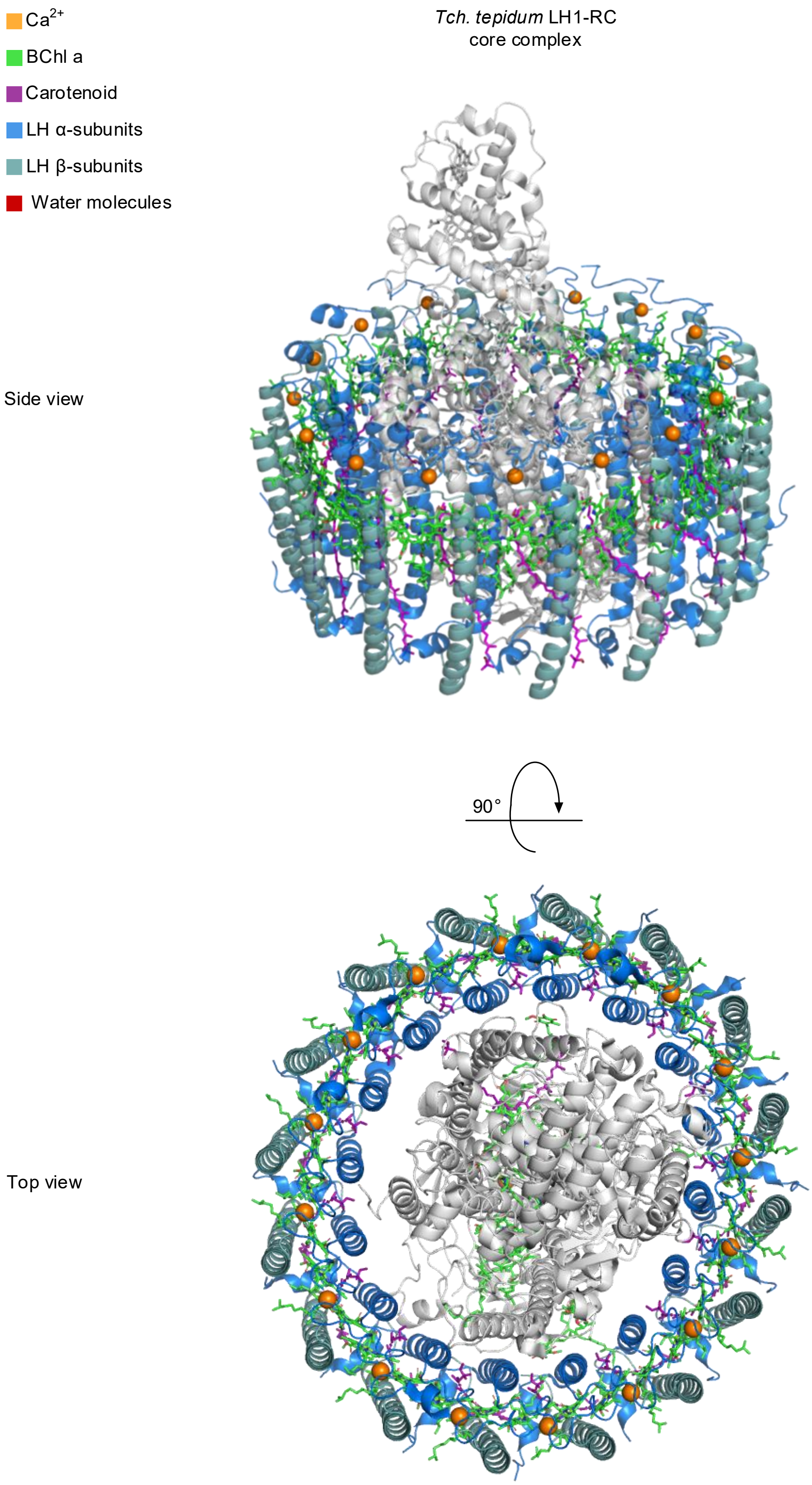


**Supplementary Figure 1.** Schematic of the overall structure of LH1-RC. The crystal structure of LH1-RC was determined at 1.9 Å resolution.

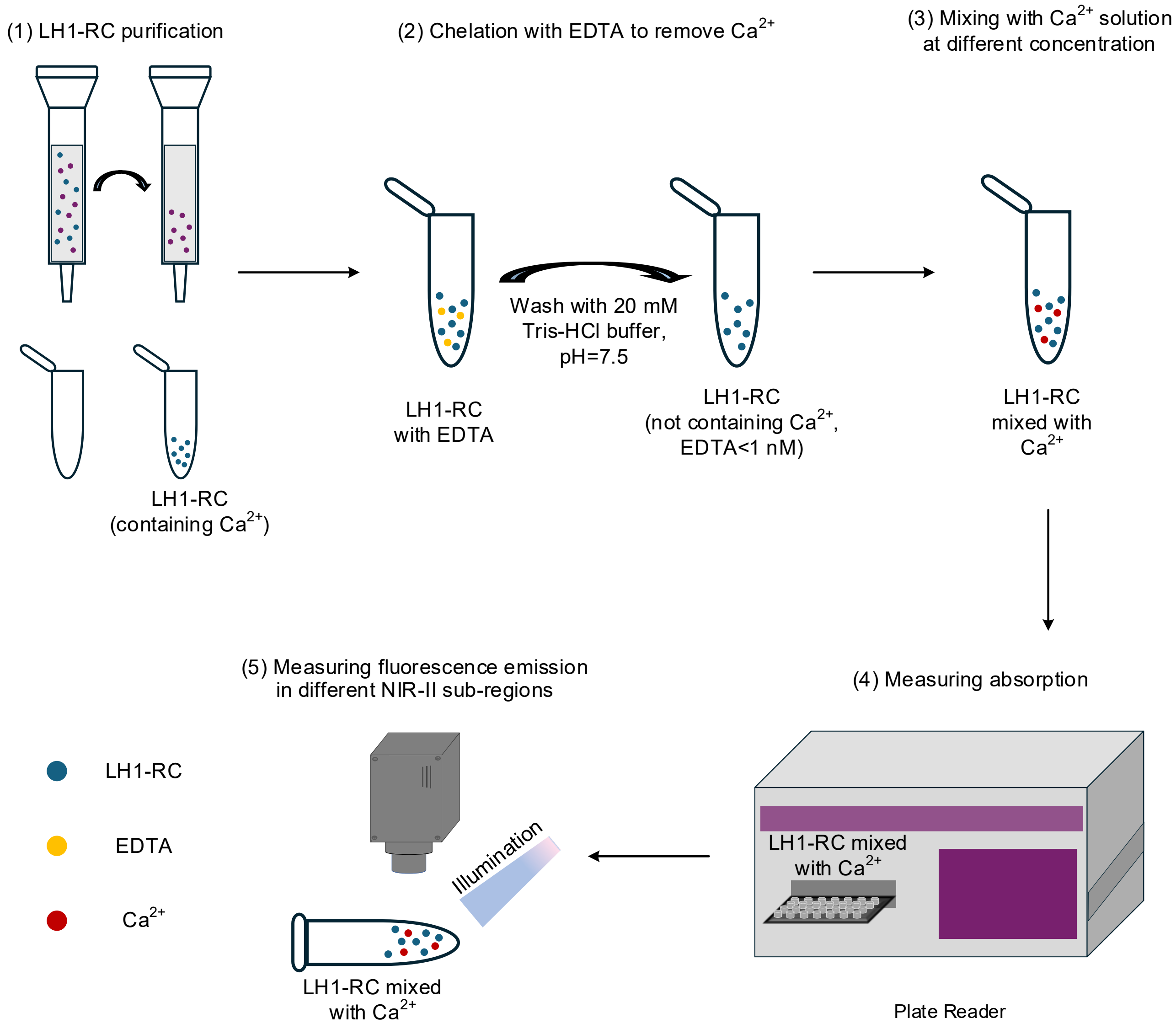


**Supplementary Figure 2. Characterization of LH1-RC.** The procedure involves (1) LH1-RC purification, (2) chelation with EDTA to remove $Ca^{2+}$, (3) mixing with $Ca^{2+}$ solution at different concentrations, (4) measuring absorption, and (5) measuring fluorescence emission in different NIR-II sub-regions.

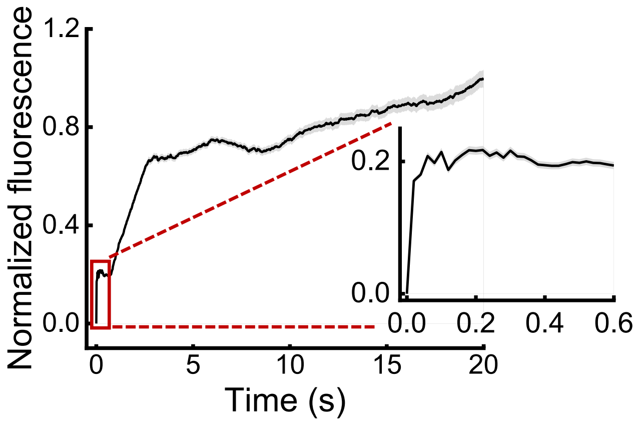


**Supplementary Figure 3.** Normalized fluorescence of LH1-RC following the addition of 10 mM $Ca^{2+}$ to 0.5 μM LH1-RC without $Ca^{2+}$. A 915 nm laser was used for excitation, and the fluorescence was filtered by a 1000 nm long-pass filter before being collected for NIR-II imaging. The exposure time was 10 ms, and the frame rate was 50 fps. Error bars represent s.d. for $n = 6$.

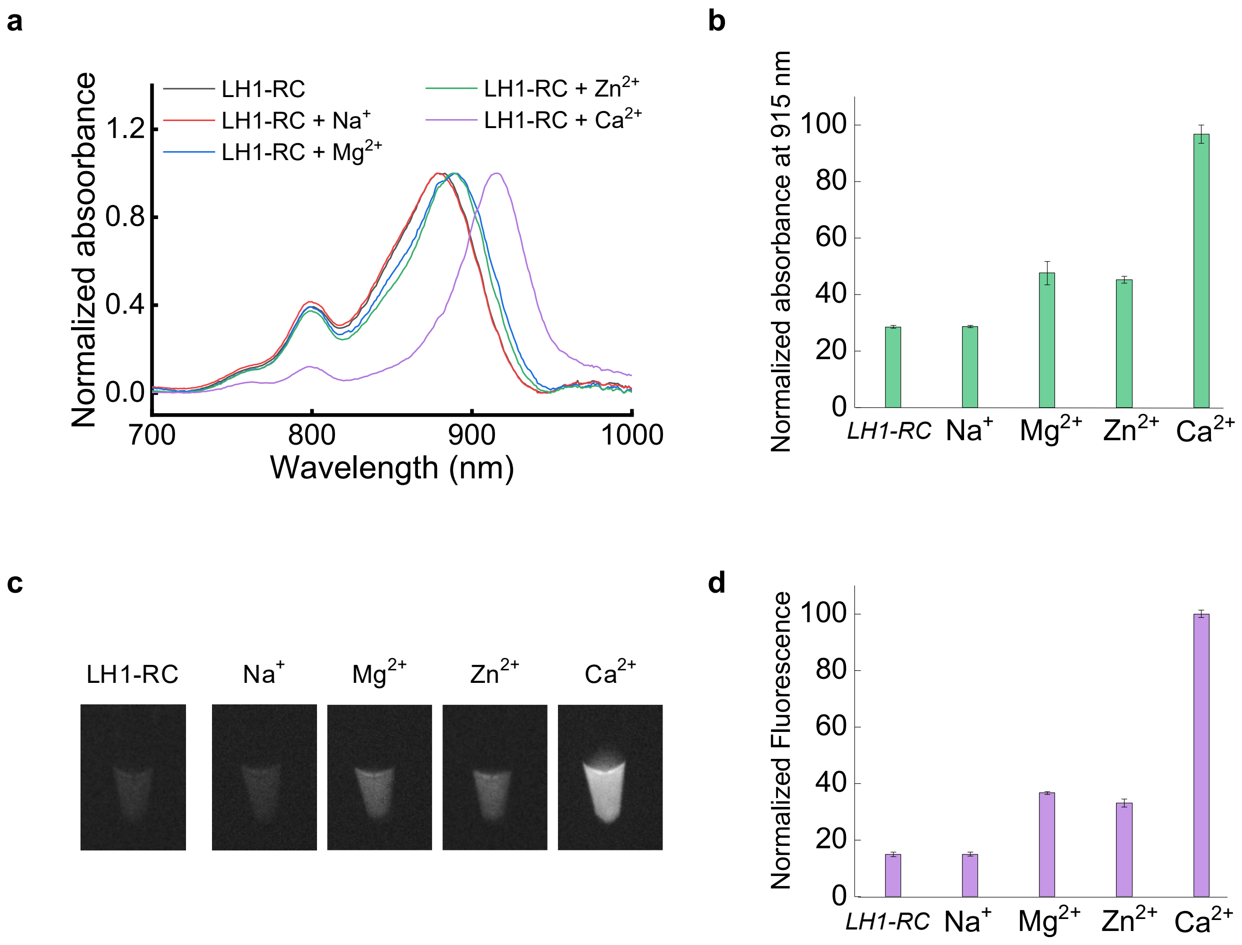


**Supplementary Figure 4. Calcium specificity analysis.** (**a**) Normalized absorbance spectra of LH1-RC before and after the addition of 1 mM $Na^{+}$, $Mg^{2+}$, $Zn^{2+}$, or $Ca^{2+}$. (**b**) Normalized absorbance of LH1-RC at 915 nm before and after the addition of 1 mM $Na^{+}$, $Mg^{2+}$, $Zn^{2+}$, or $Ca^{2+}$. All absorbance values were normalized to the absorbance of LH1-RC at 915 nm after the addition of 1 mM $Ca^{2+}$. (**c**) NIR-II fluorescence images of LH1-RC before and after the addition of 1 mM $Na^{+}$, $Mg^{2+}$, $Zn^{2+}$, or $Ca^{2+}$. A 915 nm laser was used for excitation, and the NIR-II fluorescence was filtered by a 1000 nm long-pass filter before being collected. (**d**) Normalized fluorescence of LH1-RC before and after the addition of 1 mM $Na^{+}$, $Mg^{2+}$, $Zn^{2+}$, or $Ca^{2+}$. All values were normalized to the fluorescence of LH1-RC after the addition of 1 mM $Ca^{2+}$. Error bars represent s.d. for $n = 3$.

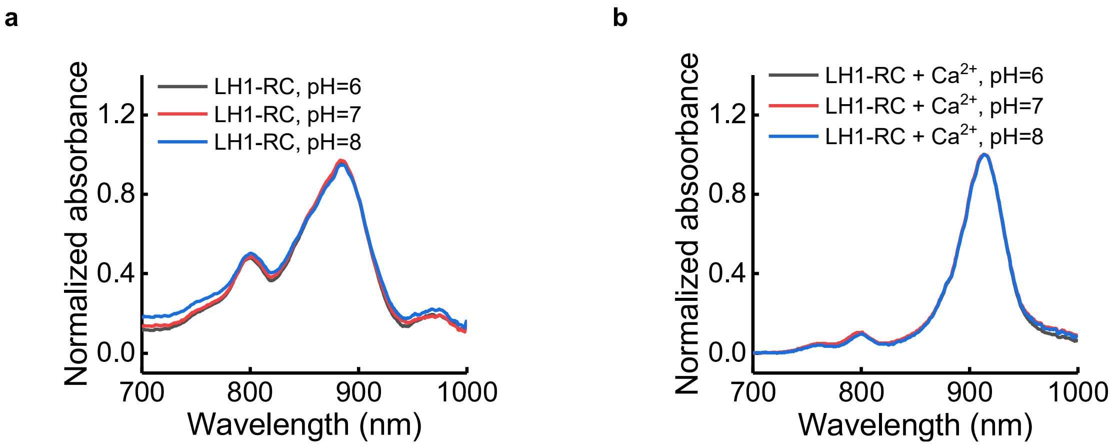


**Supplementary Figure 5.** Absorbance spectra of LH1-RC (**a**) with or (**b**) without $Ca^{2+}$ in solutions with pH 6-8.

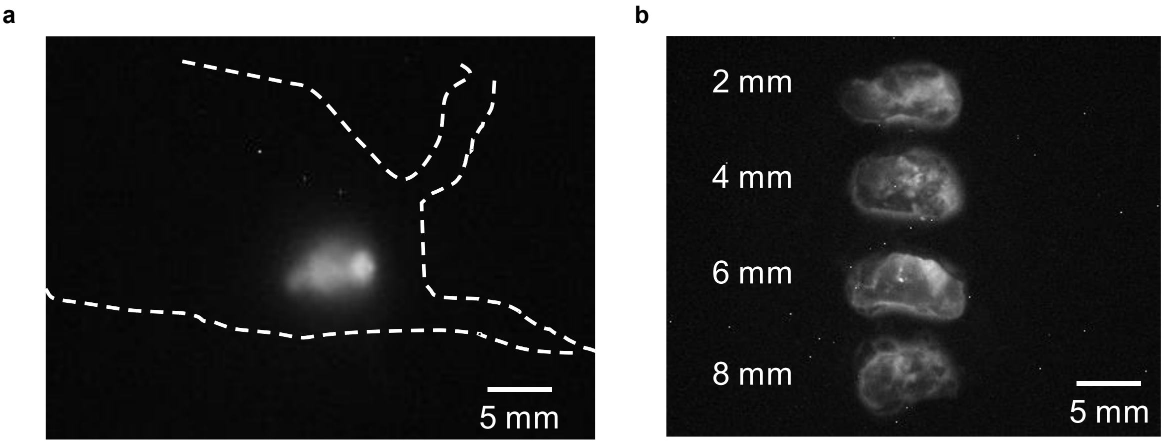


**Supplementary Figure 6.** (**a**) Wide-field imaging of a mouse bearing an A549 tumor that was injected intratumorally with LH1-RC@lipid. The exposure time was 10 ms. (**b**) A549 tumor slices at different depths after LH1-RC@lipid injection. A 915 nm laser and a 1000 nm long-pass filter were used.

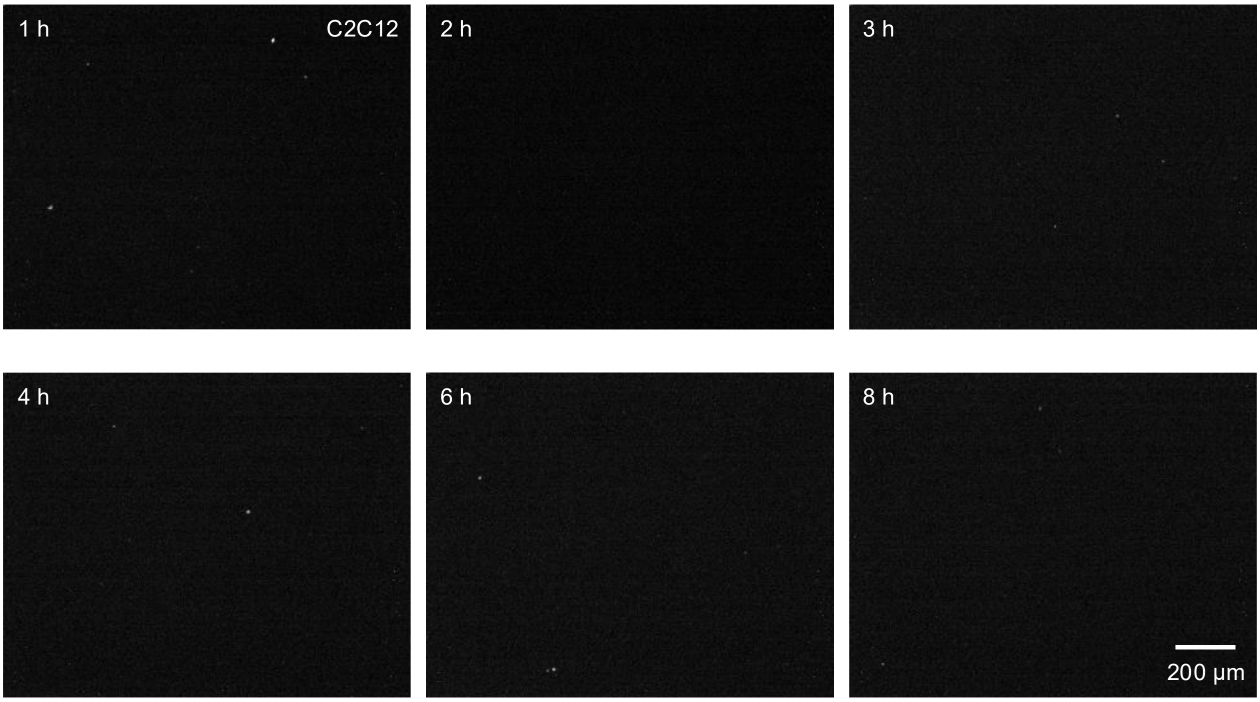


**Supplementary Figure7.** C2C12 cell uptake of pure LH1-RC at different time points after the addition of LH1-RC to the culture medium. A 915 nm laser was used for excitation, and the NIR-II fluorescence was filtered by a 1000 nm long-pass filter before being collected by an InGaAs camera. The exposure time was 200 ms. A 10× objective was used.

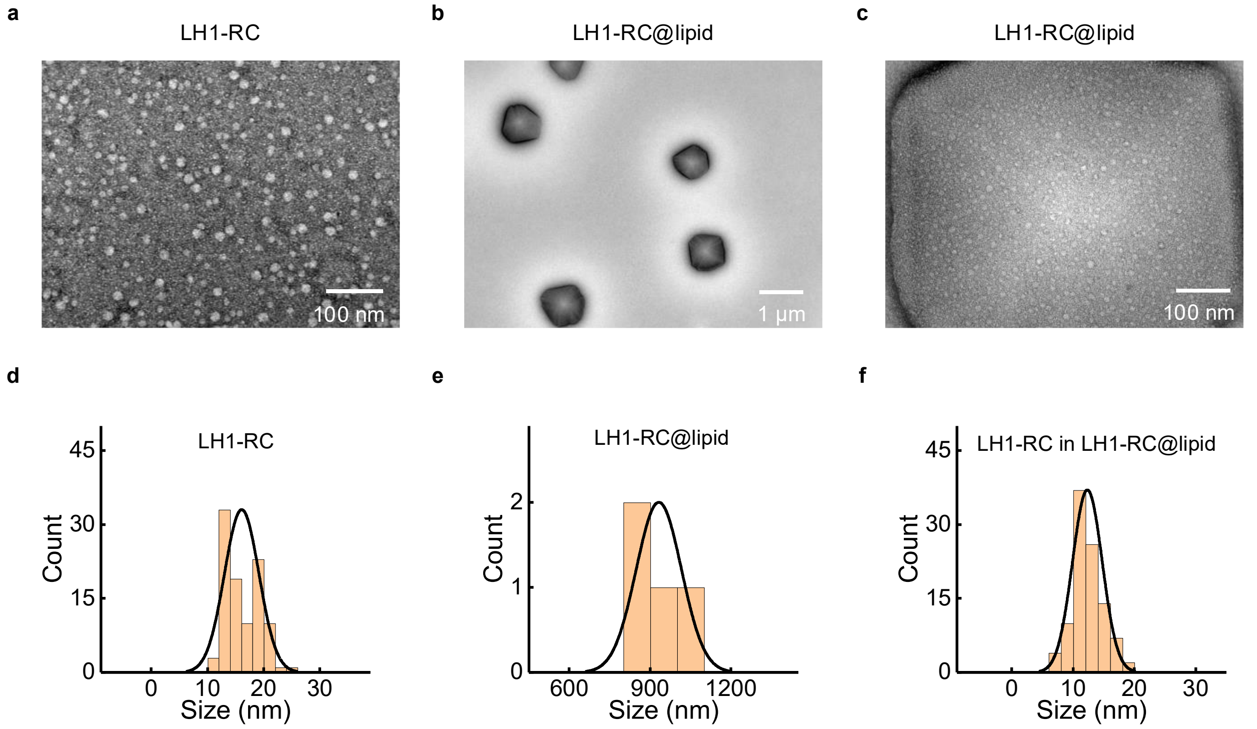


**Supplementary Figure 8.** TEM images of (**a**) LH1-RC and (**b**,**c**) LH1-RC@lipid. Size-distribution histogram of (**d**) LH1-RC ($n$ = 100, size = 16.1 ± 3.1 nm), (**e**) LH1-RC@lipid ($n$ = 4, size = 932.4 ± 83.8 nm), and (**f**) LH1-RC in LH1-RC@lipid ($n$ = 100, size = 12.3 ± 2.4 nm) measured from TEM images. The LH1-RC was encapsulated using Lipofectamine™ 3000.

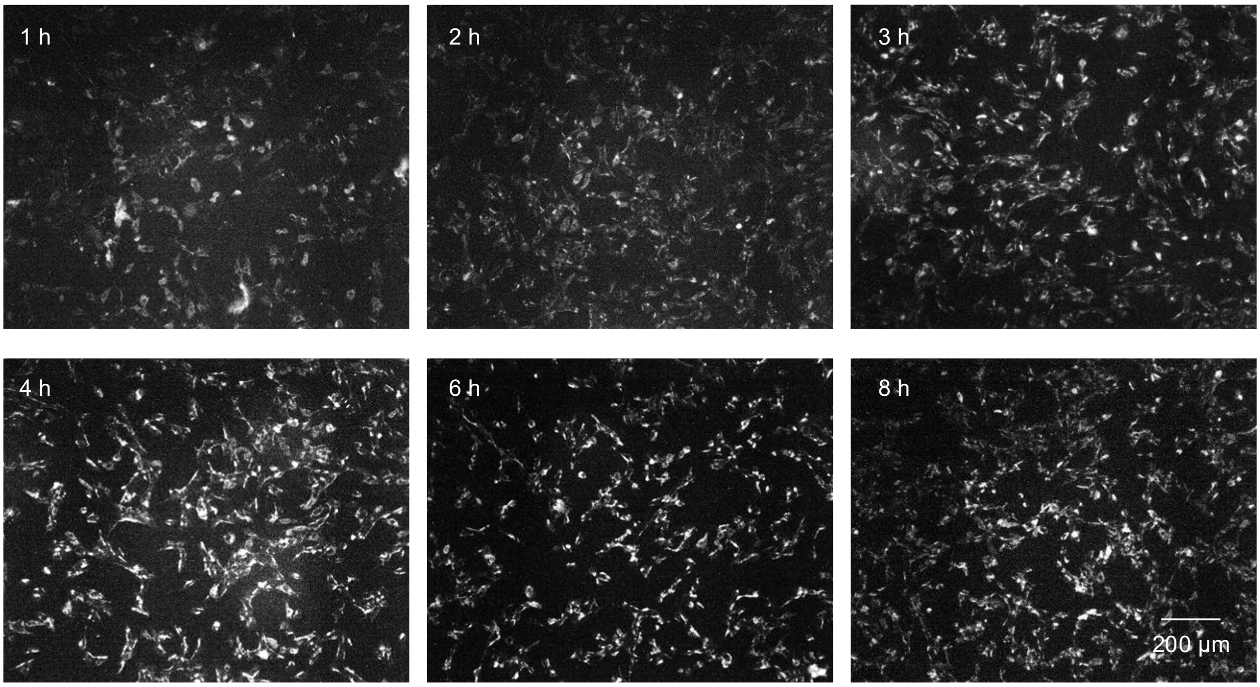


**Supplementary Figure 9.** C2C12 cell uptake of LH1-RC@lipid at different time points after the addition of LH1-RC@lipid to the culture medium. The LH1-RC was encapsulated using Lipofectamine™ 3000. A 915 nm laser was used for excitation, and the NIR-II fluorescence was filtered by a 1000 nm long-pass filter before being collected by an InGaAs camera. The exposure time was 200 ms. A 10× objective was used.

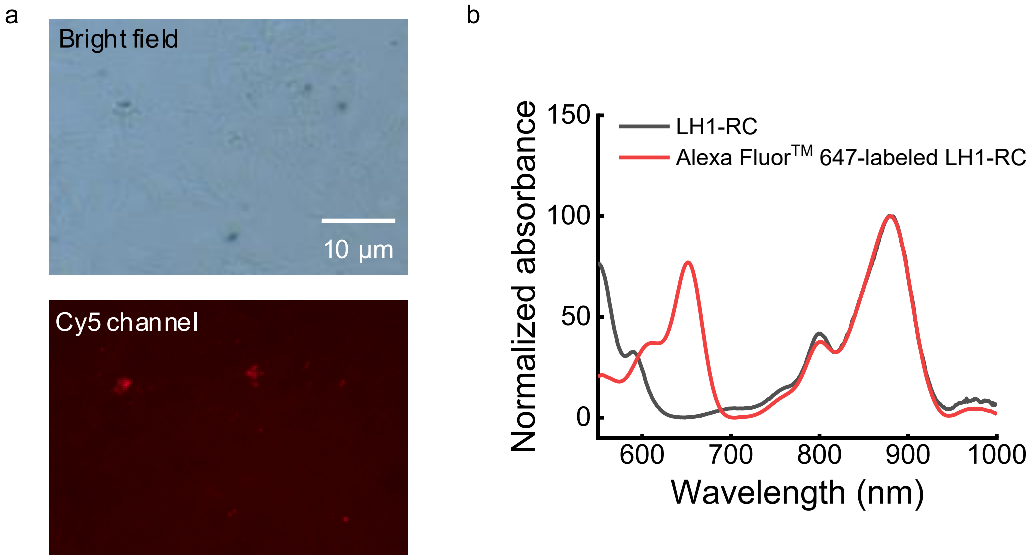


**Supplementary Figure 10.** (**a**) C2C12 cell uptake of Alexa Fluor$^{TM}$ 647@lipid at 6 h after the addition of Alexa Fluor$^{TM}$ 647@lipid to the culture medium. Alexa Fluor$^{TM}$ 647 was encapsulated using Lipofectamine™ 3000. A Cy5 filter cube and a 10× objective were used for Alexa Fluor$^{TM}$ 647 imaging. (**b**) Absorbance of LH1-RC (black) and Alexa Fluor$^{TM}$ 647-labeled LH1-RC (red), demonstrating that Alexa Fluor$^{TM}$ 647 was successfully conjugated to LH1-RC.

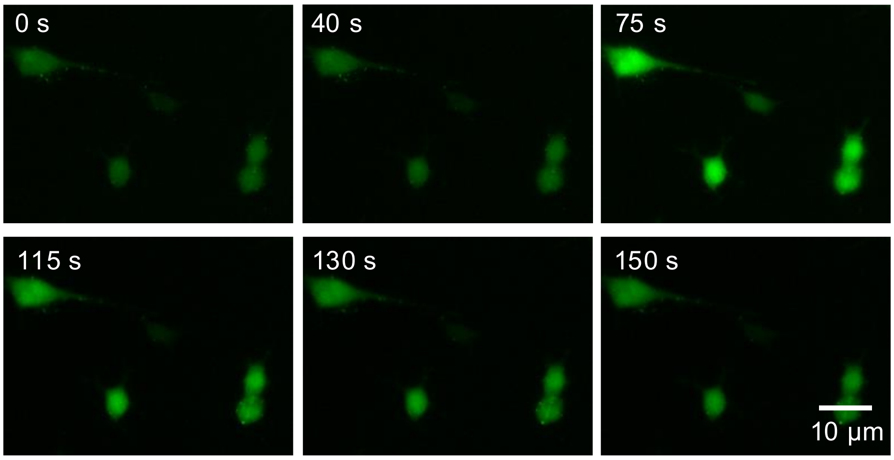


**Supplementary Figure 11**. Responses of Fluo-4 in C2C12 cells to the addition of 1 mM 4-CMC. 4-CMC was added to initiate the release of $Ca^{2+}$ from the ER/SR in C2C12 cells. A 10× objective and a FITC cube filter were used.

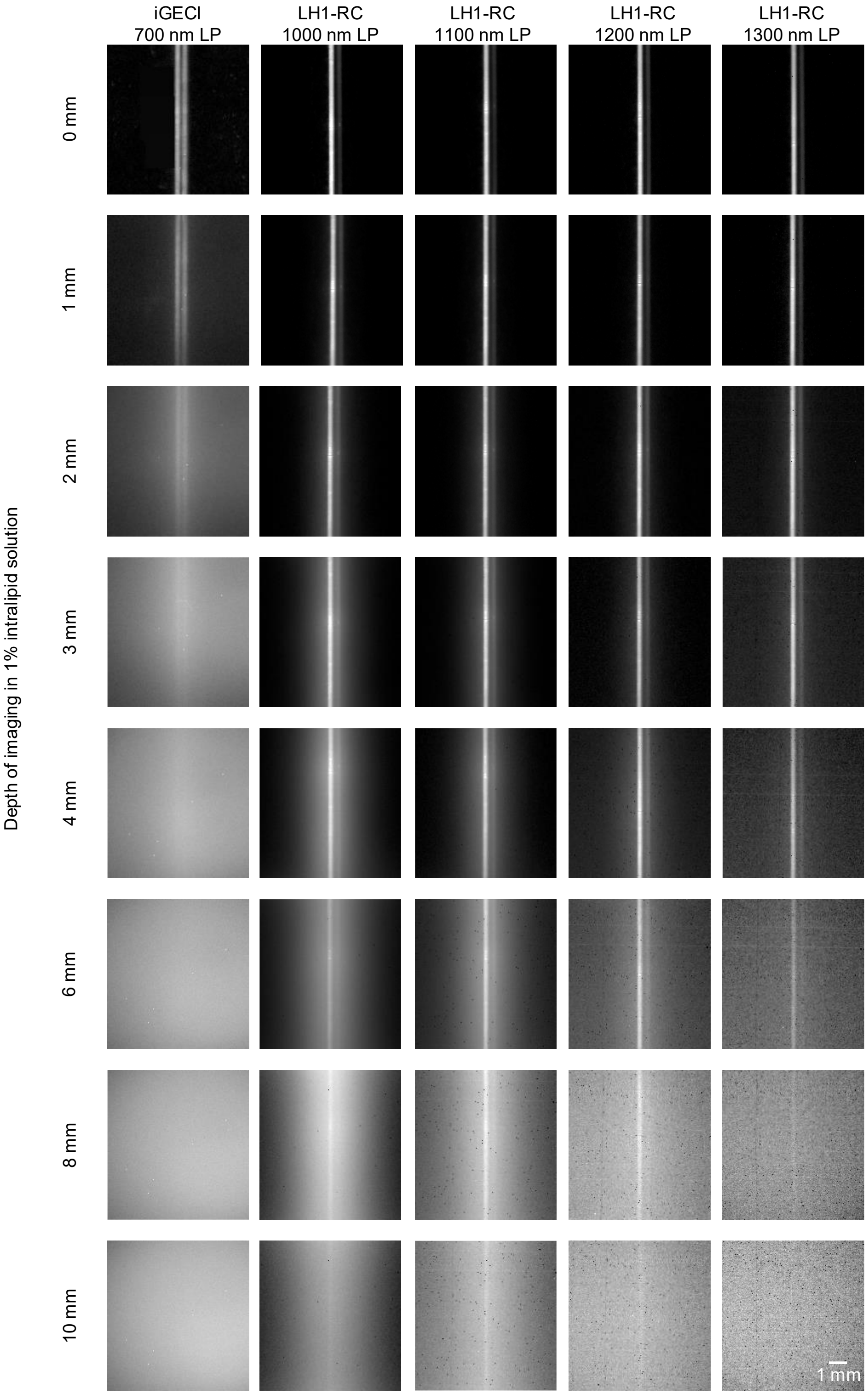


**Supplementary Figure 12. Phantom imaging in NIR-I and NIR-II sub-regions.** Fluorescence imaging of 300-μm-diameter capillary tubes filled with iGECI or LH1-RC, with or without $Ca^{2+}$, immersed at different depths in a 1% intralipid solution using a wide-field imaging system. For iGECI imaging, a 660 nm laser was used for excitation, and a 700 nm long-pass filter was applied to filter the fluorescence signal. For LH1-RC imaging, a 915 nm laser was used for excitation, and the NIR-II fluorescence was filtered using LP filters with cut-on wavelengths in the range of 1000-1300 nm.

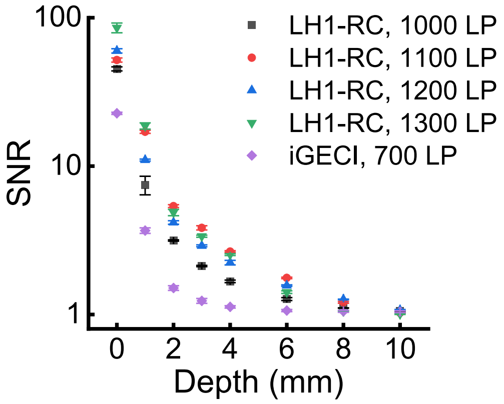


**Supplementary Figure 13.** The signal-to-noise ratios of capillaries for phantom imaging shown in Supplementary Fig. 12. Error bars represent s.d. for $n = 3$.

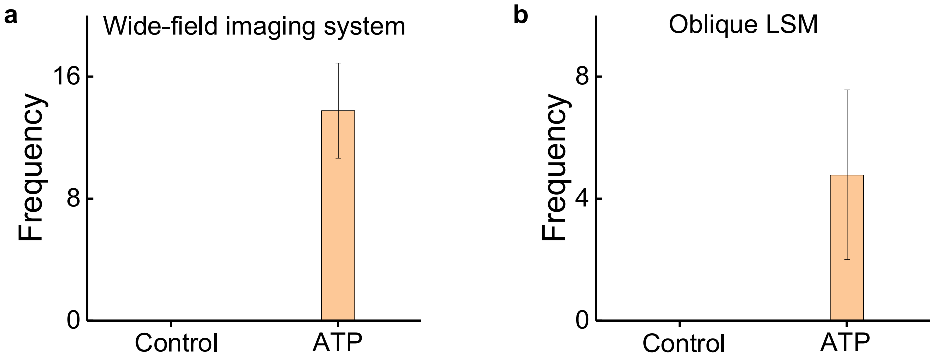


**Supplementary Figure 14.** Calcium signal frequency of HeLa tumors in response to the treatment with 1 μM ATP, as imaged by the (**a**) NIR-II wide-field imaging system or (**b**) NIR-II oblique LSM, respectively. Error bars represent s.d. for $n = 5$.

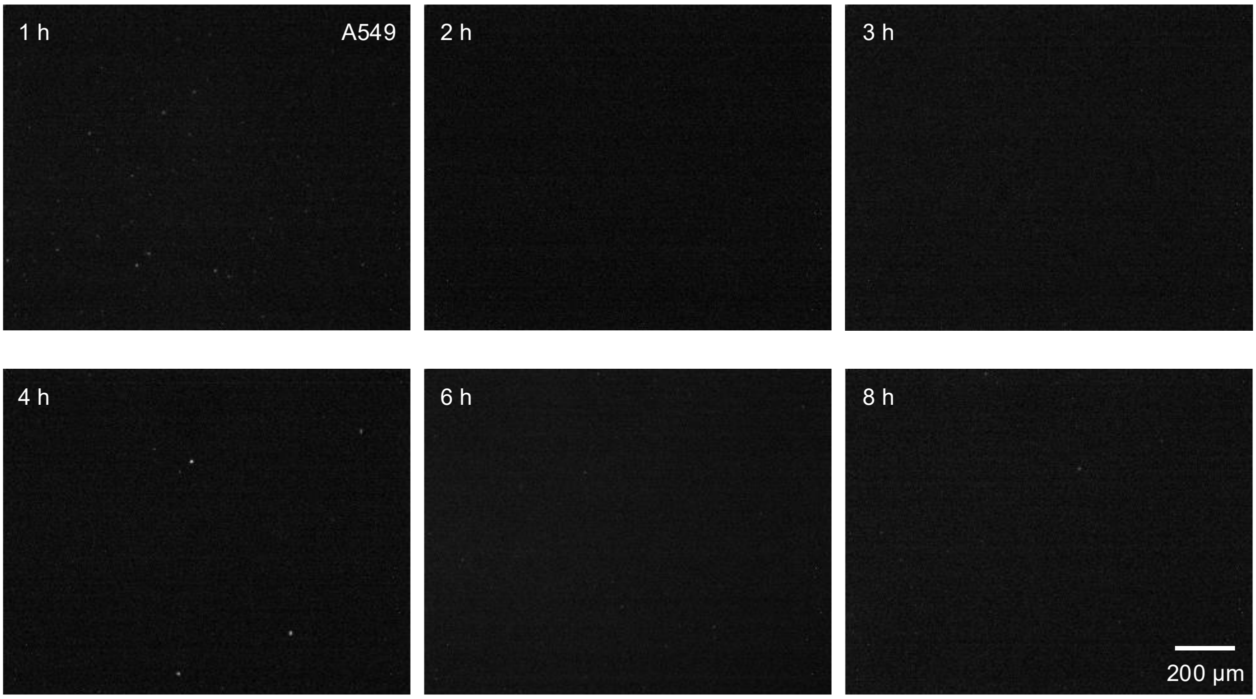


**Supplementary Figure 15.** A549 cell uptake of pure LH1-RC was assessed at different time points following the addition of LH1-RC to the culture medium. A 915 nm laser was used for excitation, and the NIR-II fluorescence was filtered by a 1000 nm long-pass filter before being collected by an InGaAs camera. The exposure time was 200 ms. A10× objective was used.

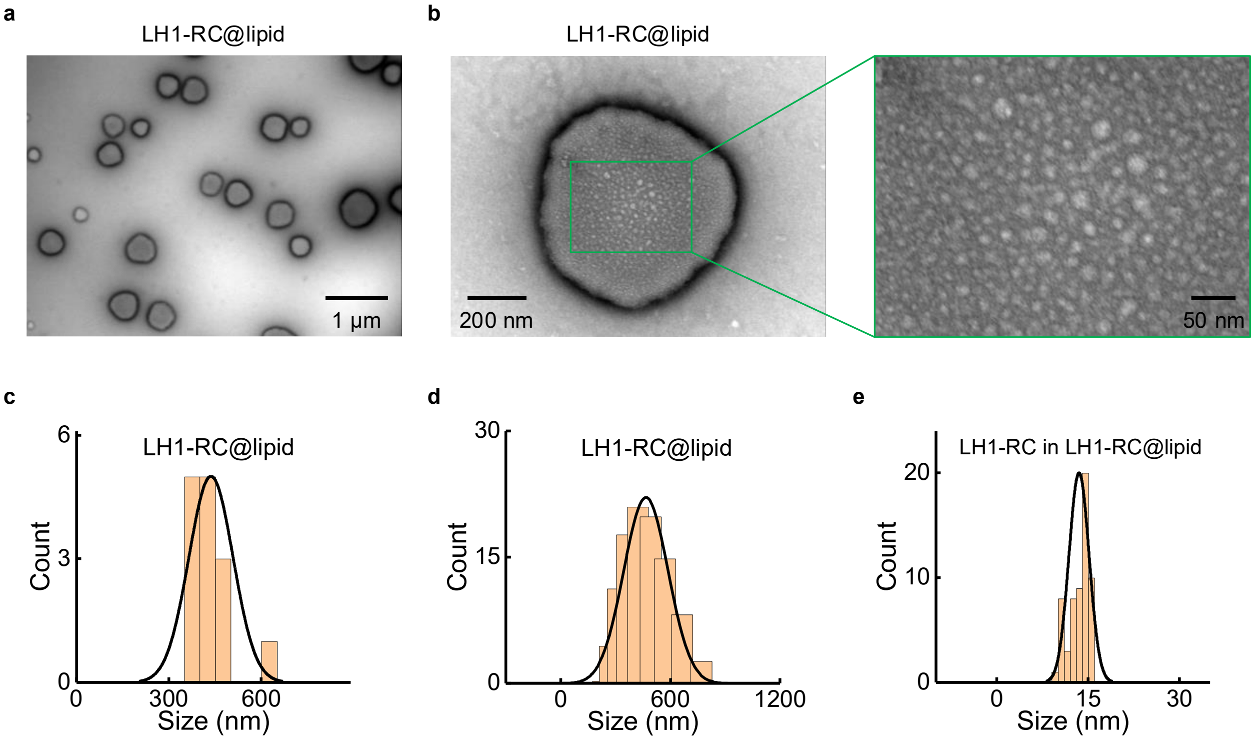


**Supplementary Figure 16.** (**a**,**b**) TEM images of LH1-RC@lipid. (**c**) Size-distribution histogram of LH1-RC@lipid ($n$ = 14, size = 436.8 ± 71.8 nm) measured from TEM images. (**d**) Dynamic light scattering spectra of LH1-RC@lipid in PBS buffer (size = 449.7 nm). (**e**) Size-distribution histogram of LH1-RC in LH1-RC@lipid ($n$ = 59, size = 13.5 ± 1.7 nm) measured from TEM images. The LH1-RC was encapsulated using Lipofectamine™ CRISPRMAX™.

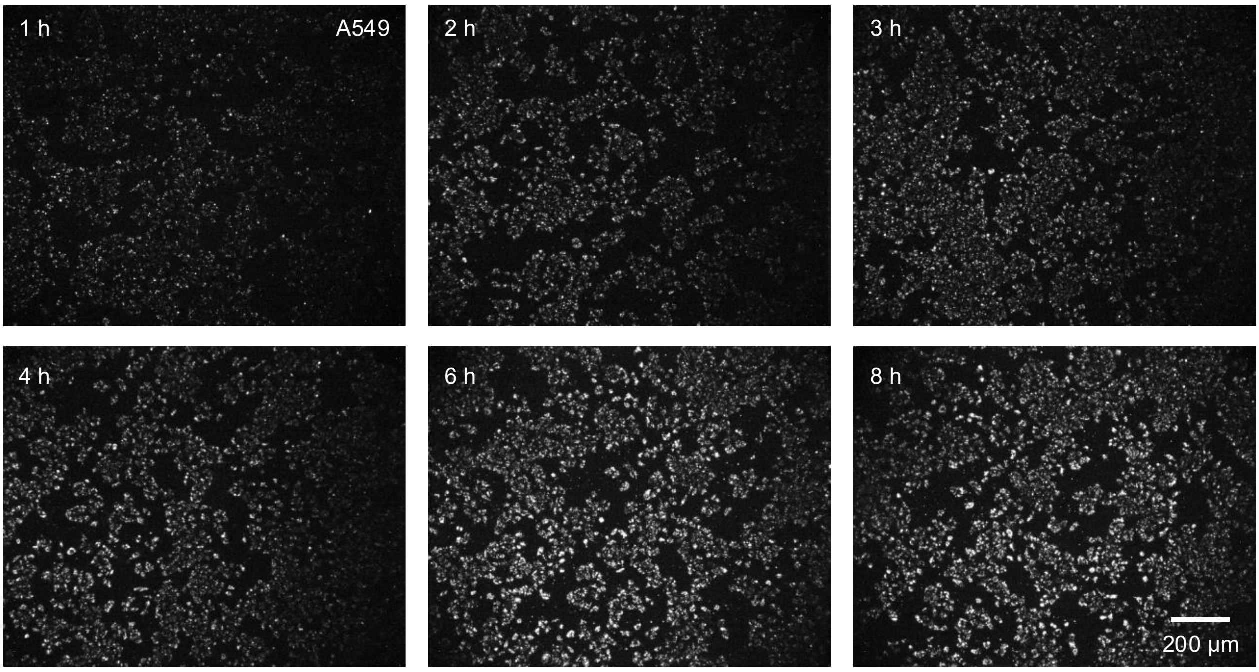


**Supplementary Figure 17.** A549 cell uptake of LH1-RC@lipid was assessed at different time points after the addition of LH1-RC@lipid to the culture medium. The LH1-RC was encapsulated using Lipofectamine™ CRISPRMAX™. A 915 nm laser was used for excitation, and the NIR-II fluorescence was filtered by a 1000 nm long-pass filter before being collected by an InGaAs camera. The exposure time was 200 ms. A 10× objective was used.

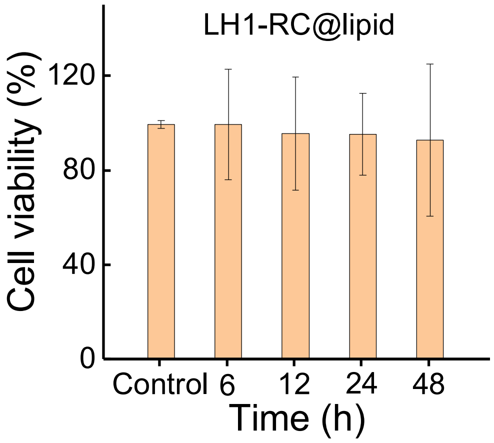


**Supplementary Figure 18.** Cell viability of A549 cells was measured after the addition of LH1-RC@lipid at different time points (6 h, 12 h, 24 h and 48 h). Error bars represent s.d. for $n = 3$.

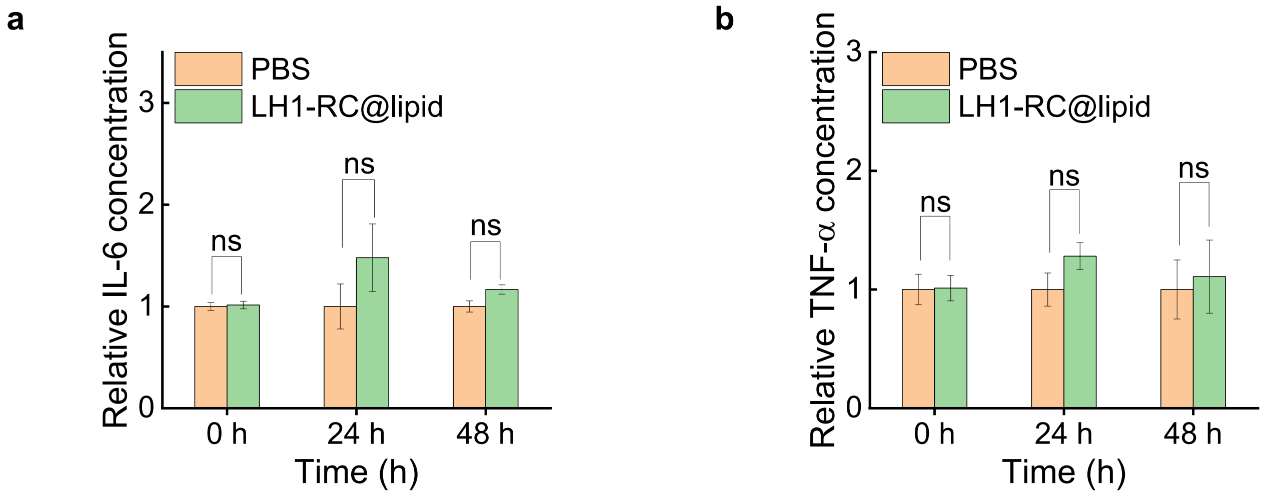


**Supplementary Figure 19.** (**a**) Relative IL-6 levels and (**b**) TNF-α levels in A549 tumor-bearing mice after injection of 50 μL of PBS or 0.03 μg/μL LH1-RC@lipid at different time points (0 h, 24 h and 48 h), as assessed by enzyme-linked immunosorbent assay (ELISA). The IL-6 and TNF-α concentrations were normalized to those of the PBS injection group at 0 h, 24 h, and 48 h, respectively. ns indicates not significant, as measured by a Pair-sample T-test.

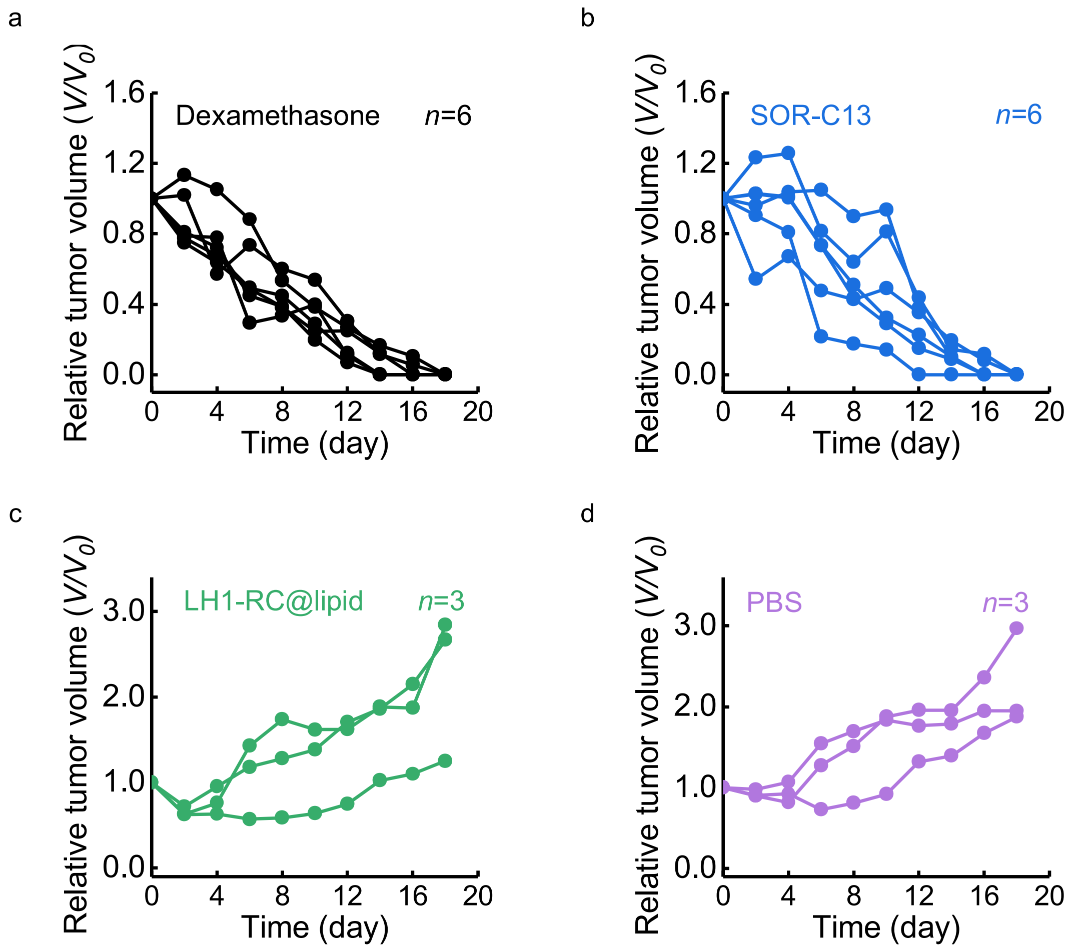


**Supplementary Figure 20.** Relative tumor volume ($V/V_0$, where $V_0$ is the tumor volume on Day 0, $V$ is the tumor volume) of A549 tumor-bearing mice was measured after treatments with (**a**) dexamethasone, (**b**) SOR-C13, (**c**) LH1-RC@lipid, and (**d**) PBS. Day 0 is defined as 4 days after tumor inoculation, with the tumor size at Day 0 being approximately 100 $mm^3$. Six mice ($n$=6) were treated with dexamethasone or SOR-C13. Three mice ($n$=3) were treated with LH1-RC@lipid or PBS.

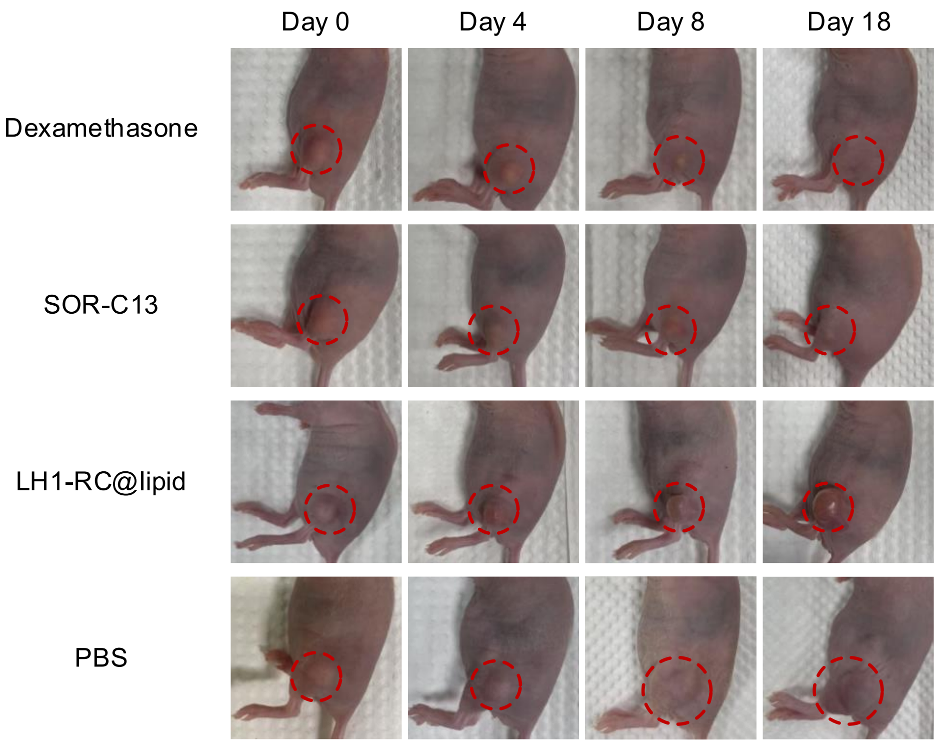


**Supplementary Figure 21.** Photographs of A549 tumor-bearing nude mice were recorded on various days post tumor inoculation (-4 days). The mice were treated on the same day (Day 0) with dexamethasone, SOR-C13, LH1-RC@lipid, or PBS.

**Supplementary Table 1. The quantum yield of LH1-RC and other NIR-II fluorophores.**

| Fluorophore | Emission (nm) | Quantum yield (QY) | Solvent | Ref. |
|---|---|---|---|---|
| IRFP1032 | 1032 | 0.84 | PBS | 1 |
| LZ1105 | 1105 | 0.03 | PBS | 1 |
| Flav7(3) | 1045 | 0.3 | Dichloromethane | 1 |
| FM1210 | 1210 | 0.03 | Dichloromethane | 1 |
| IR26 | 1138 | 0.03 | Dichloromethane | 1 |
| FBP 912 | 912 | 0.086 | Water | 2 |
| EB766 | 1530 | 0.01 | PBS | 3 |
| FT-TQT | 1034 | 0.025 | Water | 4 |
| CH-4T | 1055 | 0.06 | Water | 4 |
| CQ-4T | 1055 | 0.104 | PBS | 5 |
| dsZW1015 | 1015 | 0.014 | PBS | 6 |
| P-ipr@Gal | 1060 | 0.036 | Water | 7 |
| IR-140 | 1043 | 0.012 ± 0.007 | DMSO/Water | 8 |
| HPQ-Zzh | ~910 | 0.037 | Water | 9 |
| FD-1080/DMPC | 1370 | 0.0545 | Water | 10 |
| LH1-RC | 955 | 0.8 | Tris-HCl water | **This work** |

**Supplementary Table 2. Comparison between LH1-RC and other calcium indicators.**

| **Calcium indicators** | **Abs/Em(nm)** | **QY (%)** | **Hill coefficient** | $K_d$ | **Δ*F*/*F* (Purified protein)** | **Δ*F*/*F* (*In vivo*,mouse)** | **Ref.** |
|---|---|---|---|---|---|---|---|
| jGCaMP8s | 488/510 | 53 | 2.2 ± 0.1 | 46 ± 1 nM | 49.5 ± 0.1 | / | 11 |
| G-CatchER+ | 488/510 | 64 ± 1 | / | 1.2 ± 0.2 mM | 0.6 | / | 12 |
| GCaMP6m | 497/515 | 61 | 3 | 167 nM | / | 13% (1 AP, cell-attached recording) | 13 |
| HaloGFP-Ca1–4 | 498/513 | 72 | / | 9.2 µM | 9.2 | / | 14 |
| GreenT-EC | 504/515 | 72 | 0.8 ± 0.1 | 7.1 ± 1.2 mM | 61 | / | 15 |
| HaloCaMP1a-$JF_{614}$ | 628/646 | 74 | / | 892 nM | 29.5 | / | 16 |
| CaSiR-2 AM | 636/661 | 26 | / | 0.31 µM | / | / | 17 |
| iGECI | donor 640/670 acceptor 700/720 | / | 2.5 and 0.9 | 15 and 890 nM | 6 | 23% (Wide-field imaging through cranial window) | 18 |
| Ca-NIR | 666/684 | 0.6 | / | 8 µM | / | / | 19 |
| NIR-II Ca | 677/935 | / | / | / | / | / | 20 |
| NIR-GECO1 | 678/704 | 1.9 | 1.03 | 215 nM | -9 | 0.3% (Wide-field imaging through cranial window) | 21 |
| WHaloCaMP | 679/690 | 50 | 2.5 ± 0.3 | 37 ± 2 nM | / | ~25% (Two-photon imaging through cranial window) | 22 |
| $2XR_{715}$ -HaloCaMP1a | 725/760 | / | / | 45 nM | 1.7 | / | 23 |
| LH1-RC | 915/ >1000 | 0.8 | 2.7 | 63.7 µM | 6.2 | 3.6% (Non-invasive wide-field imaging) | **This work** |

Notes: Abs is absorbance wavelength; Em is emission wavelength; QY is quantum yield; $K_d$ is affinity constant; AP is action potential.

**Supplementary Table 3. The comparison between LH1-RC-based one-photon NIR-II imaging and two- or multi-photon imaging using visible-wavelength $Ca^{2+}$ indicators.**

| Calcium indicators | Em (nm)/ Ex(nm) | Imaging depth (µm) | FOV (µm$^2$) | Frame rate (fps) (pixels) | SNR | Imaging type | Invasive | Ref. |
|---|---|---|---|---|---|---|---|---|
| GCaMP6s | 515/ 920 | 250 | 400 × 400 | 5 (512 × 512) | / | 2PM (*in vivo*, cortex) | Yes | 24 |
| Fluo-4 acetoxymethyl ester (AM) | 520/ 800-880 | 200-300 | 600 × 600 | 1.95 (512 × 256) | / | 2PM (*in vivo*, cortex) | Yes | 25 |
| GCaMp6s | 515/ 1300 | 978 | 250 × 250 | 8.35 (200 × 200) | / | 3PM (*in vivo*, hippocampus) | Yes | 26 |
| GCaMP6s | 515/ 1300 | 1000 | 250 × 250 | 8 (256 × 272) | / | 3PM (*in vivo*, cortex) | Yes | 27 |
| GCaMP6s | 515/ 1300 | 984 | 200 × 200 | 8.49 (256 × 256) | >40 | 3PM (*in vivo*, hippocampus) | Yes | 28 |
| Fluo-4 acetoxymethyl ester (AM) | 519/ 495 | / | 360 × 360 | 0.5 ( / ) | / | One-photon wide-field imaging (*in vitro*, tumor slice) | / | 29 |
| Fluo-4 acetoxymethyl ester (AM) | 519/ 495 | 4.8 | 606.1 × 606.1 | 0.5 (512 × 512) | / | CM (*in vitro*, tumor slice) | / | 30 |
| LH1-RC | >1000/ 915 | 8000 | 5000 × 5000 | 50 (640 × 512) | 45.2 | One-photon wide-field | No | **This** |

| | | | | | | | | |
|---|---|---|---|---|---|---|---|---|
| | | | | | | imaging (*in vivo*, tumor) | | **wo rk** |

Notes: Em and Ex are emission and excitation wavelengths, respectively. FOV: field of view; SNR: signal-to-noise ratio; 2PM: two-photon fluorescence microscopy; 3PM: three-photon fluorescence microscopy; CM: confocal microscopy.
